# Phosphorylation-induced flexibility of proto-oncogenic Bcl3 regulates transcriptional activation by NF-κB p52 homodimer

**DOI:** 10.1101/2024.06.30.601400

**Authors:** Wenfei Pan, Tapan Biswas, Shandy Shahabi, William Suryajaya, Andres Vasquez, Qian Du, Gourisankar Ghosh, Vivien Ya-fan Wang

## Abstract

B cell lymphoma 3 (Bcl3), a member of the IκB family proteins, modulates transcription by primarily associating with NF-κB p50 and p52 homodimers. Bcl3 undergoes extensive phosphorylation, though the functions of many of these modifications remain unclear. We previously described that phosphorylation at Ser33, Ser114 and Ser446 partially switches Bcl3 from acting as an IκB-like inhibitor to a transcription regulator by associating with the (p52:p52):DNA binary complex. Here, we identified another critical phosphorylation site, Ser366. Substituting at all four residues to phospho-mimetic glutamate further enhances Bcl3’s transcriptional activity. Phospho-modifications retain Bcl3’s ability to stably bind p52 but induces reciprocal structural changes as revealed by HDX-MS experiments; the N-terminal region stiffens, while the C-terminus becomes more flexible. The increased flexibility allowed the Bcl3:(p52p52) binary complex to better accommodate DNA. The removal of the C-terminal 28-residues transformed Bcl3 into a transcriptional activator independent of phosphorylation. Notably, most identified mutations in Bcl3 from various cancers map to its C-terminus, suggesting the functional relevance of Bcl3 C-terminal structural flexibility and enhanced interaction with (p52p52):DNA complex to transcriptional potential and disease. Overall, this study uncovers the mechanistic basis by which phosphorylation-driven structural changes convert Bcl3 from an inhibitor to a transcriptional cofactor of NF-κB, and how deregulation of its activity through altered phosphorylation or mutation can lead to cancer.

## Introduction

B-cell lymphoma 3 (Bcl3) was initially discovered due to its constitutive expression in many B cell cancers (Ohno et al. 1993), and later observed to be highly expressed in various cancers, including solid tumors, due to either aberrant chromosomal translocation of the gene or overexpression without chromosomal rearrangement (Nishikori et al. 2003; Mathas et al. 2005; Maldonado and Melendez-Zajgla 2011). Bcl3 belongs to the inhibitor of the NF-κB (IκB) family of proteins that regulate the transcriptional activities of the NF-κB family of transcription factors. Three classical members of the IκB family, IκBα, -β and -ε, form binary complexes with homo- and hetero-dimers of RelA and cRel of the NF-κB family in the cytoplasm to prevent their access to and association with DNA in the nucleus, thereby inhibiting their transcriptional activities (Basak et al. 2007; Huxford et al. 2011). In contrast, Bcl3 associates with only the p52:p52 and p50:p50 homodimers of NF-κB family in the nucleus (Bours et al. 1993; Fujita et al. 1993; Inoue et al. 1993). Bcl3 associates with pre-formed p50:p50 homodimer while it induces the formation of p52:p52 homodimer in cells although the heterodimeric complexes of p52 with RelA, RelB and cRel are more stable (Pan et al. 2022). IκBζ, another IκB family protein, functions similarly to Bcl3 as transcriptional coregulator by preferentially associating with p50:p50 homodimer in the nucleus (Bates and Miyamoto 2004; Yamamoto et al. 2004; Trinh et al. 2008; Kohda et al. 2016).

The biological function of Bcl3 is diverse; however, its most notable function is the induction of cell proliferation by activating the expression of cyclin D1 (Cogswell et al. 2000; Westerheide et al. 2001). Bcl3 also facilitates the development of splenic B cells, the expansion and recall response of memory T cells, and the priming of CD4 and CD8 cells by dendrites (Franzoso et al. 1997; Schwarz et al. 1997). Deletion of the Bcl3 gene does not show any overt phenotype in cells; however, it causes poor immune response when these cells are challenged with infection. This observation tallies with a protective function of the lack of Bcl3 in autoimmune diseases such as colitis and experimental autoimmune encephalomyelitis (EAE) (Liu et al. 2022). The mechanism underlying function of Bcl3 as a transcriptional coregulator or its preference for only two specific NF-κB dimers, p52:p52 and p50:p50 homodimers, remains elusive till date. Tissue-specific variation in expressions of p50 and p52, and their alternative production mechanisms suggest that functions of Bcl3 is likely to be cell type- and signal-specific.

The structural organization of Bcl3 is similar to other IκB proteins, containing a centrally folded ankyrin repeat domain (ARD) flanked by flexible N- and C-terminal regions (Fig. 1A-B). Prototypical IκB proteins (IκBα, -β and -ε) are rarely involved in transcriptional activity in the nucleus. Rather, their ubiquitination and proteosomal degradation upon phosphorylation at the N-termini by signal-induced IκB kinase (IKK) leads to nuclear translocation of NF-κB dimers and transcriptional activation. Bcl3 is not reported to have such phosphorylation sites regulating its degradation. Instead, phosphorylation of serine 33 (S33) located at the N-terminal region of Bcl3 is critical for its nuclear localization, and consequent protection from constitutive cytoplasmic degradation (Wang et al. 2017). Bcl3, IκBα and -β all contain proline and serine rich (PEST) sequences in their C-termini. Phosphorylation at the PEST sequence in IκBα renders it a stronger inhibitor (Phelps et al. 2000) but modification of these serine residues to asparagine render a transcriptional coactivator function to IκBα (Dembinski et al. 2017). In contrast, Bcl3 is heavily phosphorylated at its C-terminal serine and threonine residues in cells, some of the phosphorylation being critical for transcriptional activation (Wang et al. 2017). The primary focus of this study is to clarify mechanisms underlying the phosphorylation-mediated transcriptional coregulatory functions of Bcl3.

**Figure 1.**
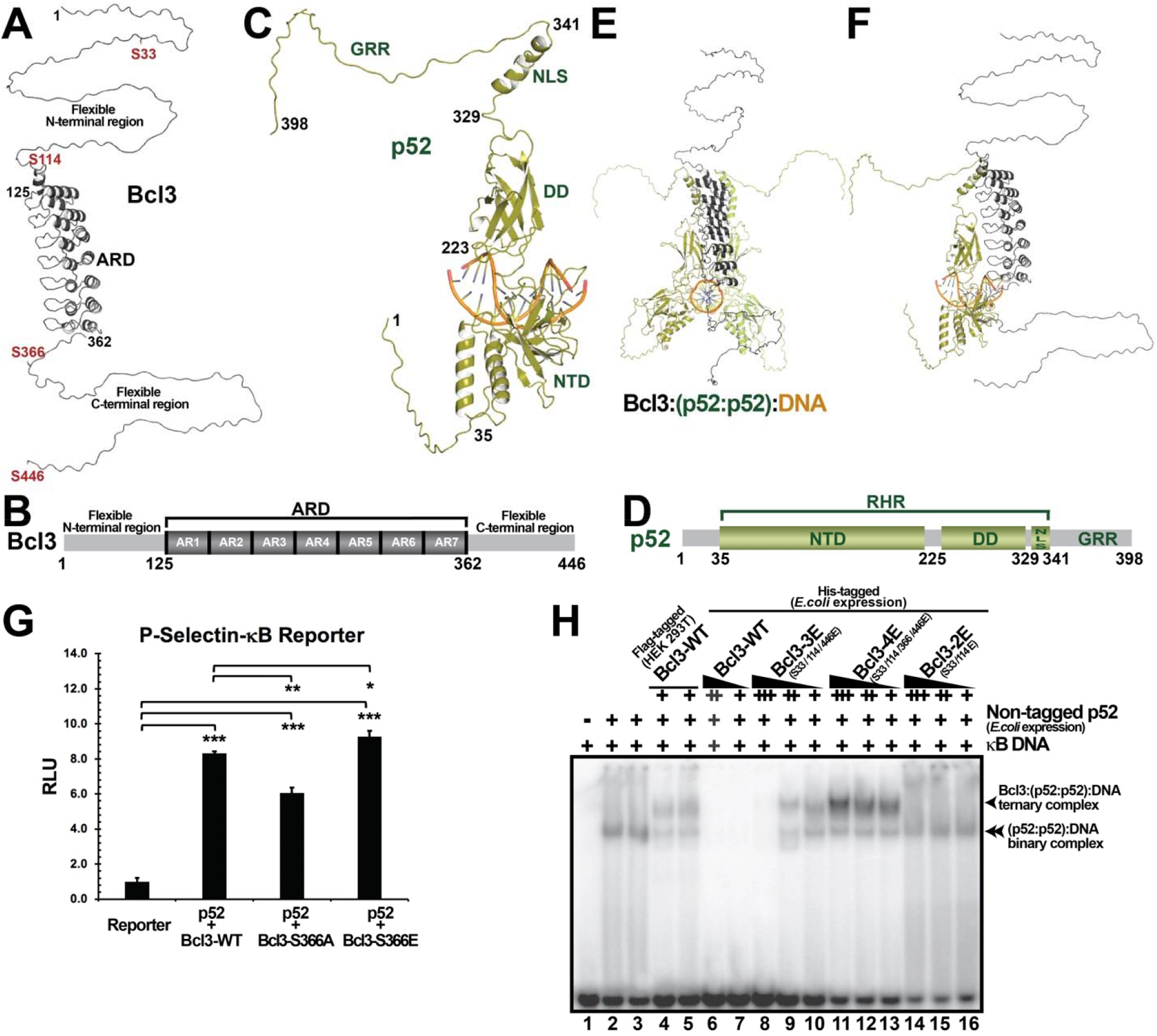
Phosphorylation of Bcl3 at S366 is critical for transcriptional activation by p52. (A, B) Structural organization of Bcl3 (the sequence numbering corresponds to UniProt ID P20749 lacking eight N-terminal residues) indicating ankyrin repeat domain (ARD) bounded by flexible N- and C-terminal regions; positions of serine residues undergoing phosphorylation investigated here are indicated in red. (C, D) Structural organization of p52 protomer (residues 1-398 of 900 from UniProt ID Q00653) observed in context of a homodimer in association with a DNA-duplex; the residues at the boundaries of N-terminal domain (NTD), dimerization domain (DD), nuclear localization signal (NLS) and glycine rich region (GRR) boundaries are indicated. (E, F) Orthogonal views of predicted structural mode of Bcl3, p52:p52 homodimer and DNA ternary complex; side view in (F) is showing only one protomer of p52 for clarity. (G) Luciferase reporter activity of a κB-promoter driven by co-expression of p52 and Bcl3. The S366A substitution of Bcl3 reduced the reporter activity while the S366E substitution enhanced it. The data were analyzed from three independent experiments performed in triplicate. RLU, relative luciferase unit. *p<0.05; **p<0.01; ***p<0.001 (t test). Error bars represent standard deviation (SD). (H) EMSA of κB-DNA with p52:p52 homodimer (expressed in *E. coli*) in absence or presence of Flag-tagged Bcl3 (expressed in HEK 293T cells, thus likely phosphorylated) respectively displaying formation of (p52:p52):DNA binary complex (lanes 2-3) and Bcl3:(p52:p52):DNA ternary complex (lanes 4-5). Expulsion of DNA from the (p52:p52):DNA binary complex by Bcl3-WT expressed in *E. coli* thus unphosphorylated (lanes 6-7). Formation of an unstable ternary complex by (p52:p52):DNA with Bcl3-3E^(S33/114/446E)^ expressed in *E. coli* (lanes 8-10), and at a higher concentration Bcl3-3E competes with DNA for binding to p52:p52 homodimer dissolving the ternary complex (lane 8). Formation of a stabler ternary complex by (p52:p52):DNA with Bcl3-4E^(S33/114/366E446E)^ expressed in *E. coli*, which is perturbed only weakly at a high concentration of Bcl3-4E (lanes 11-13). Absence of a ternary complex formation by (p52:p52):DNA with Bcl3-2E^(S33/114E)^ expressed in *E. coli*, and corresponding lack of ability to displace DNA from (p52:p52):DNA by Bcl3-2E (lanes 14-16).

The bacterial or baculovirus expressed Bcl3 (presumably unphosphorylated) fails to form a ternary complex with (p52:p52):DNA (Nolan et al. 1993); rather it expels DNA from p52:p52 homodimer to form a stable Bcl3:(p52:p52) binary complex. A structural model of the Bcl3:(p52:p52):DNA ternary complex, predicted from structural knowledge of other NF-κB:IκB and NF-κB:DNA subcomplexes, indicates a clash of the C-terminal region and PEST sequence of Bcl3 with the negatively charged DNA (Fig. 1E-F). This agrees with the stripping of DNA from homodimers of p50 and p52 by unphosphorylated Bcl3 (Nolan et al. 1993) (Fig. 1H, lanes 6-7). This stripping effect of Bcl3, which reflects its functional resemblance to inhibitory IκBα, is not yet reported in cells. Earlier, we observed that phosphorylation at S33 of Bcl3 is required for its nuclear localization, and that at S114 and S446 for the formation of the Bcl3:(p52:p52):DNA ternary complex (Wang et al. 2017). S114 is located within the flexible N-terminal region just next to the ARD, and S446 is the C-terminal residue of Bcl3. Phospho-inactive mutation of S114 and S446 of Bcl3 to alanine abolished cyclin D1 expression and inhibited the migratory function of U2OS cells. However, if expressed and isolated from mammalian cells, this mutant protein could still form a ternary complex with p52 and DNA (Wang et al. 2017). This hints at more complex possibilities through various post-translational modifications of Bcl3 at other site(s), some of which will enable Bcl3 to form a ternary complex, and perhaps trigger transcriptional activation of specific genes.

In this study, we observed that the additional phosphorylation of Bcl3 at S366 plays a critical role in promoting the formation of a stable Bcl3:(p52p52):DNA ternary complex and activating the transcriptional of a reporter gene in cells. Addition of this new phosphorylation site converts Bcl3 having two phosphorylation sites on either side of the central ARD spanning residues 126 to 362. To probe the effect of phosphorylation on structural properties of Bcl3 that help stabilize the Bcl3:(p52p52):DNA ternary complex for regulation of transcriptional activation, we performed hydrogen-deuterium exchange mass spectrometry (HDX-MS) experiments on seven recombinant proteins or complexes, p52 homodimer, Bcl3-WT, Bcl3-4E, Bcl3-WT:(p52:p52), Bcl3-4E:(p52:p52), (p52:p52):DNA and Bcl3-4E:(p52:p52):DNA. The deuterium uptake data revealed key interaction features of p52 and Bcl3, as well as phosphorylation-induced structural changes in Bcl3. Our findings demonstrate that the phosphorylation drives extrusion of Bcl3 C-terminal region from the DNA-binding interface of p52:p52 homodimer for the accommodation of cognate DNA. The resulting ternary complex is weaker than the Bcl3-4E:(p52:p52) binary complex, since contacts disrupted by the C-terminal extrusion are only partially compensated by some new interactions formed between p52 and Bcl3-4E away from the protein-DNA interface. The truncation of the C-terminal 28-residues of Bcl3 recapitulated phosphorylation-driven functional switch. The significance of this region is underscored by several oncogenic mutations, including frameshifts, as noted in the COSMIC database (https://cancer.sanger.ac.uk/cosmic/gene/analysis?ln=BCL3) (Forbes et al. 2015). These findings highlight the importance of the C-terminal domain in phosphorylation-dependent activity switch of Bcl3 and its importance in transcriptional regulation.

## Results

### Phosphorylation of S366 in Bcl3 is essential for its association with the promoter and transcriptional activation

We set out to determine if other phosphorylation, in addition to phosphorylation at S33, S114 and S446 of Bcl3 could enhance its ability to associate with the (p52:p52):DNA binary complex and induce gene activation. Examination of mass-spectrometry data from our previous experiments and publicly available proteomic databases of different cancer cells identified new phosphorylation sites that have not been studied. We identified phosphorylation at S366 in our sample and observed that it is common in cancer cells. S366 is located at the C-terminal flexible region of human Bcl3 which is in the vicinity of the DNA-binding interface of p52 in the ternary complex model (Fig. 1A-B, 1E-F). Hence, we investigated if phosphorylation at S366 could play a regulatory role. Bcl3 is constitutively phosphorylated when over-expressed in cells. Therefore, phospho-inactive S336A and phospho-mimetic S366E mutants were generated to study the effect of phosphorylation at S366. Using a reporter gene expression assay, we observed a reduction in p52-dependent transcriptional activation upon S366A substitution, whereas a slight enhancement upon S366E substitution comparing to the constitutive phosphorylated Bcl3-WT in cells (Fig. 1G). The difference in transcriptional output between these two mutants is the result of the phosphorylation state at S366, although it is important to note that these phospho-inactive and phospho-mimetic Bcl3-S366A and Bcl3-S366E proteins are likely to have undergone common phospho-modifications at other sites such as at S33, S114 and S446 in cells.

To test the effect of the S366E modification on the stability of the Bcl3:(p52:p52):DNA ternary complex, we performed electrophoretic mobility shift assays (EMSA) with WT Bcl3 and its multiple variants using bacterial recombinant proteins which allow us to specifically access the effect on these mutants since no other sites are subjected to modification in bacteria; the ternary complex formed with Bcl3-4E was stable at a higher concentration (Fig. 1H, lanes 11-13). The co-elution of Bcl3-4E, p52, and DNA observed in size-exclusion chromatography further supports the increased stability of the Bcl3-4E:(p52:p52):DNA ternary complex (Supplemental Fig. S1A-F).

### Impact of phosphorylation on Bcl3 dynamics in absence of p52

Structural information about the N- and C-terminal regions where phosphorylatable residues (S33, S114, S366 and S446) under study here is lacking likely due to their flexible nature (Fig. 1A-B). To explore the dynamics of the N-terminal (1-125), ARD (126-362), and C-terminal (363-446) regions as well as interactions among them, we performed HDX-MS analyses (Oganesyan et al. 2018; Ramirez-Sarmiento and Komives 2018; Vinciauskaite and Masson 2022) to measure deuterium uptakes at 0.5-, 1-, 2-, and 5-minute timepoint on Bcl3-WT and Bcl3-4E as free proteins (Fig. 2A-C). For free Bcl3-WT, 107 distinct peptides were identified with an average redundancy of 4.04, and this rendered a coverage of 98% of the protein with only two short segments spanning residues 1-6 and 249-251 missing (Supplemental Fig. S2). For Bcl3-4E, a total of 88 distinct peptides were identified with an average redundancy of 3.16, providing 99% coverage with only a small segment spanning residues 249-251 missing. The deuterium uptake plots reveal the most significant difference between these two molecules lies in the 9-45 peptide (Fig. 2A, 2D; Supplemental Fig. S5). The deuterium uptake of this peptide is negligeable in Bcl3-4E compared to Bcl3-WT suggesting the rigidity of this region in Bcl3-4E. This segment contains S33E which is known to stabilize Bcl3 from degradation by preventing the binding of ubiquitin E3 ligase TBLR1 (Keutgens et al. 2010). The structural stability of this segment explains why TBLR1 fails to bind the S33 phosphorylated Bcl3 allowing it to move to the nucleus for its transcriptional activity.

**Figure 2.**
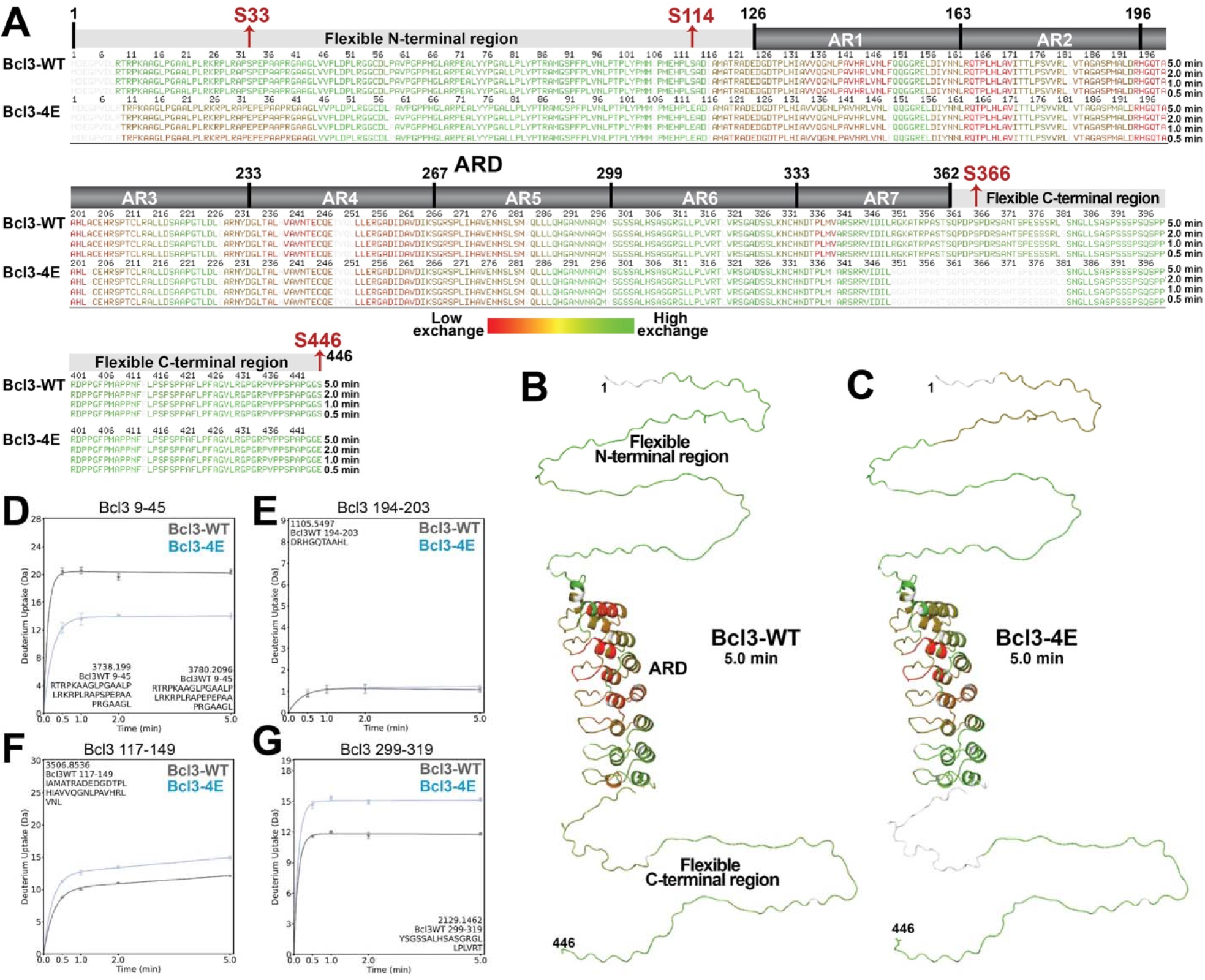
Dynamics of free Bcl3 protein. Deuterium uptake efficiencies of Bcl3-WT and Bcl3-4E as free proteins (A) mapped on their sequences at 0.5-, 1-, 2-, and 5-min timepoints; and (B, C) displayed on their structural models generated using experimentally determined structure of Bcl3-ARD and hypothetical extension of flexible regions at 5-min timepoint, respectively using red to green color spectra (red indicates low exchange, and green indicates high exchange). The N- and C-terminal domains of Bcl3 show higher exchange (reflecting flexibility) relative to the individual ARs within the ARD domain (reflecting structural rigidity). (D) Superposed deuterium uptake plots of a peptide (residues 9-45) displaying a contrasting difference in exchangeability between Bcl3-WT (high exchange in grey color) and Bcl3-4E (low exchange in blue color). The deuterium uptake plots of representative Bcl3-WT (grey) and Bcl3-4E (blue) peptides with (E) slow (residues 194-203), (F) intermediate (residues 117-149), and (G) fast (residues 299-319) exchanges, respectively. All analyses were performed in experimental triplicate with error bar shown.

Except for the 9-45 region, the deuterium uptake pattern is highly similar between the two proteins. However, the extent of protection is higher in Bcl3-WT (exchanged less) than Bcl3-4E (Fig. 2A). The N- and C-terminal regions of both Bcl3-WT and Bcl3-4E proteins displayed high exchange, suggesting their significant flexibility; however, a peptide encompassing residues 9-45 is more resistant to exchange in Bcl3-4E compared to Bcl3-WT as mentioned above. Since S33E substitution is located within this segment, local structural change induced by S33E most likely results in this resistance. The exchange propensity of individual ankyrin repeats (ARs) within the ARD displayed striking variability for Bcl3-WT and Bcl3-4E. As anticipated for a folded domain, AR1-5 is resistant to exchange with AR2 and AR3 manifesting the highest resistance (Fig. 2E). However, the deuterium uptakes of AR6 and AR7 are noticeably higher, indicating their flexibility and reflecting their structural adaptability to accommodate DNA (Fig. 2G). Noticeably, stretches of both AR1 (141-148) and AR7 (329-339) were significantly more flexible in Bcl3-4E than Bcl3-WT (Fig. 2A, 2F-G). It is possible that substitutions of S114E and S366E in nearby regions led to these effects in AR1 and AR7 of Bcl3-4E, respectively. Overall, these HDX-MS data on deuterium uptake propensity reveal that barring a segment within the N-terminal region surrounding S33E which is more rigid, the rest of Bcl3-4E is more flexible than Bcl3-WT, suggesting a role of phospho-mimetic substitutions in regulating the activity of Bcl3 through dynamism.

### Dynamics of p52 in the absence of DNA

To observe dynamics of p52 in its homodimeric complex, we performed HDX-MS analyses to measure deuterium uptakes at 0.5-, 1-, 2-, and 5-minute timepoints on p52 within its dimer (Fig. 3). Like all other NF-κB family protein, p52 has a highly conserved N-terminus known as the Rel homology region (RHR) which can be folded into the N-terminal domain (NTD), dimerization domain (DD) and nuclear localization signal (NLS) (Fig. 1C-D). The DD mediates NF-κB proteins homo- and hetero-dimerization; the NTD and DD together are responsible for DNA binding; the NLS region is flexible in solution and together with the DD forms the binding sites for IκB proteins. p52 also has a short glycine rich region (GRR) at its C-terminus. As anticipated, the N-terminal segment (1-34) and the C-terminal GRR region (351-398) exchanged deuterium rapidly suggesting their flexibility (Fig. 3A-B). The NTD showed mixed properties, with some areas exchanging rapidly and some highly resistant (Fig. 3C-3E). The DD was observed to be rather resistant to exchange, indicating its stabler fold (Fig. 3A; Supplemental Fig. S3). In addition, we also measured the uptakes of p52:p52 homodimer in the presence of its cognate DNA and/or Bcl3. The exchange propensity of various regions changed upon binding to DNA and/or Bcl3. These changes are described in detail in a later section.

**Figure 3.**
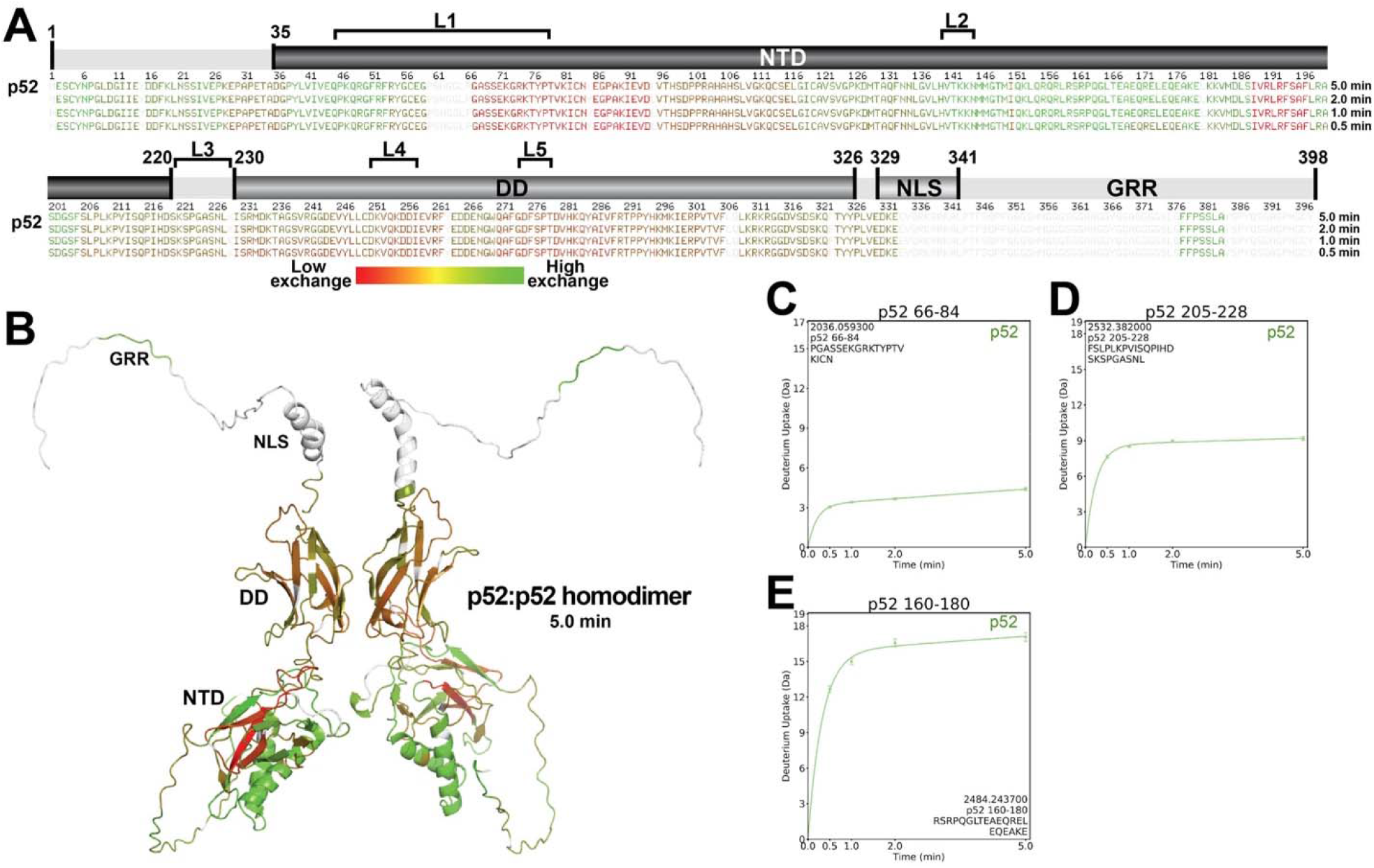
Dynamics of free p52 homodimer. Deuterium uptake efficiencies of free p52 (A) mapped on its sequence at 0.5-, 1-, 2-, and 5-min timepoints; and (B) displayed on its structural model of p52:p52 homodimer adapted from its experimentally determined structure and hypothetical extension of N- and C-terminal flexible regions at 5-min timepoint, respectively using red to green color spectra (red indicates low exchange, and green indicates high exchange). The protomers of p52 are colored identically based on average exchange of the two protomers in the homodimer; however, differential orientation and depth of the protomers in the 3D-projected view does not reflect this symmetry optimally. The deuterium uptake plots of representative p52 peptides with (C) slow (residues 66-84), (D) intermediate (resides 205-228), and (E) fast (residues 160-180) exchanges, respectively. All analyses were performed in experimental triplicate with error bar shown.

### Changes in dynamics of Bcl3 within Bcl3:(p52:p52) binary complex and Bcl3:(p52:p52):DNA ternary complex

We further performed HDX-MS studies of the p52 homodimer bound to Bcl3-WT and Bcl3-4E to understand how phospho-mimetic mutations of Bcl3 alter its binding dynamics with p52. We also performed this study using Bcl3-4E bound to both p52 homodimer as a binary complex and (p52:p52):DNA as a ternary complex to better understand how the phospho-mimetic changes help accommodate DNA. The HDX-MS data indicated unique dynamism in Bcl3 upon its binding to p52 homodimer or (p52:p52):DNA complex. One of the striking features of these complexes is that the differences in exchange between free Bcl3 and p52-bound Bcl3 are very similar in both Bcl3-WT vs. Bcl3-4E binary complexes where they showed similar exchange patterns of protection or exposure, but Bcl3-4E is more dynamic both in free and p52-bound states (Fig. 4A; Supplemental Fig. S5). That is, Bcl3-WT is more protected with less deuterium exchanges than Bcl3-4E in both states (Fig. 4F-4J). In addition to uptake plots of Bcl3 in the complexes, we also plotted differences in exchanges in the binary complexes of the two Bcl3 proteins (Bcl3-WT:(p52:p52) – Bcl3-WT, and Bcl3-4E:(p52:p52) – Bcl3-4E) which show the relative effect of p52 on these two Bcl3 proteins (Fig. 4B-C). As indicated earlier, the N-terminal segment, residues 9-45, is protected in free Bcl3-4E (Fig. 2A, 2G) but exposed in the binary complex with p52 (Fig. 4C, 4E); however, the dynamicity of this region in Bcl3-4E changes further significantly from free in Bcl3-4E:(p52:p52) binary complex to rigid in Bcl3-4E:(p52:p52):DNA ternary complex (Fig. 4C-D). The corresponding region in Bcl3-WT does not change upon binding to p52 (Fig. 4B). The rest of the N-terminal region in Bcl3-WT, residues 46-115, showed no significant difference in exchange in free and p52-bound states (Fig. 4B). This corresponding segment in Bcl3-4E is slightly more protected in the Bcl3-4E:(p52:p52) binary complex than in its free state (Fig. 4C). Most of AR1 (126-159) was protected in both binary complexes; the entire AR2-4 (160-266) is slightly exposed in both binary complexes; and the last three ARs (AR5-7, 267-362) are more protected in both cases. The C-terminal region of Bcl3-WT do not show many changes, while Bcl3-4E is slightly protected by p52.

**Figure 4.**
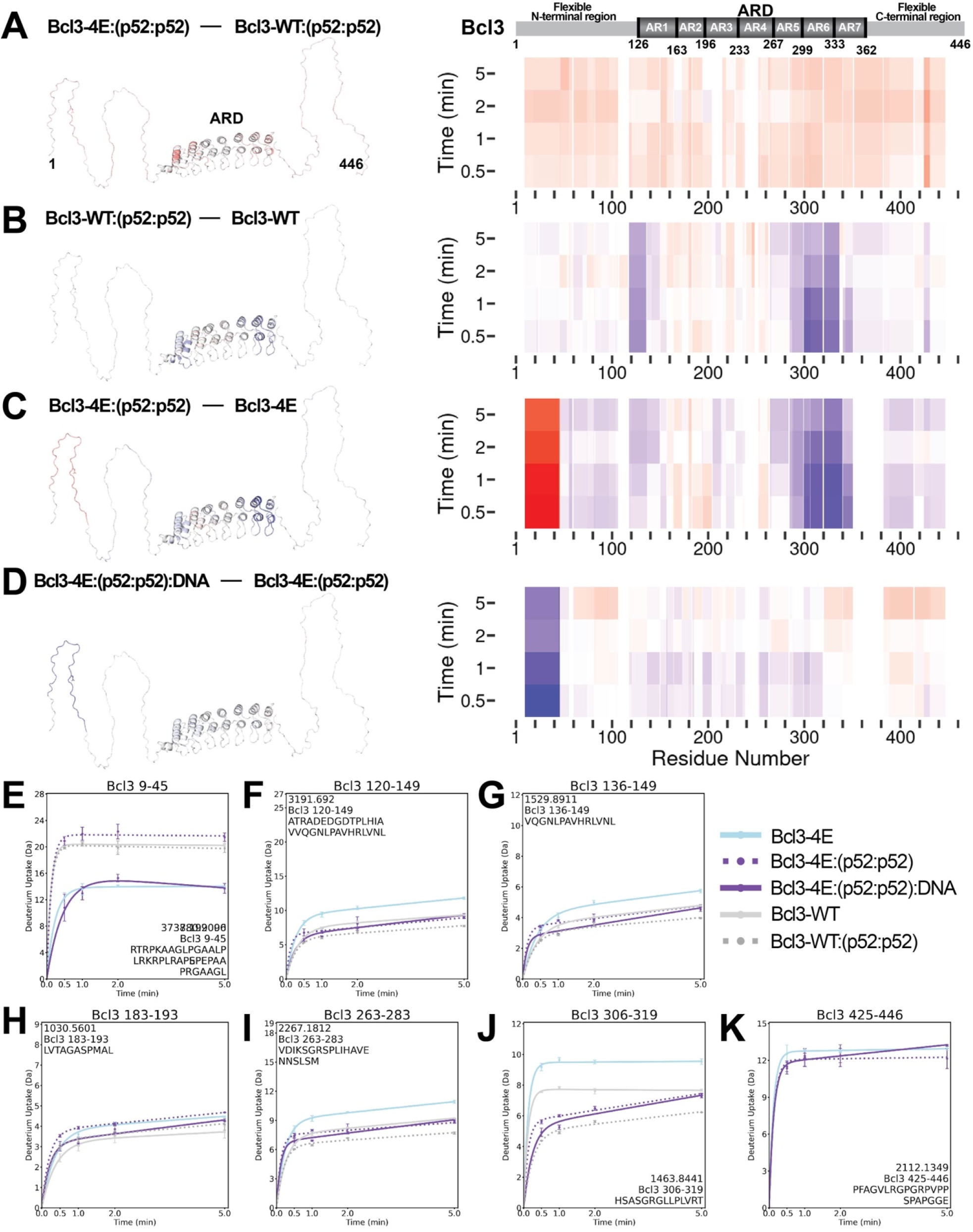
Dynamics changes of Bcl3 upon binding to p52 homodimer and (p52:p52):DNA complex. The difference in fractional uptake of deuterium in (A) Bcl3-4E vs. Bcl3-WT in their (p52:p52)-bound states, (B) Bcl3-WT in (p52:p52)-bound vs. unbound states; (C) Bcl3-4E in (p52:p52)-bound vs. unbound states; and (D) Bcl3-4E in (p52:p52):DNA-bound vs. (p52:p52)-bound states. The differences in fraction uptake displayed on Bcl3 structural model are shown on the left; sequence ascribed heatmaps are shown on the right. Changes are represented in Blue-White-Red spectra (with grey representing absence of data, blue signifies protection, and red signifies deprotection), the indicated range of which is based on the spread of the differences (Supplemental Fig. S4). The deuterium uptake plots of representative Bcl3 peptides (E) residues 9-45; (F) residues 120-149; (G) residues 136-149; (H) residues 183-193; (I) residues 263-283; (J) residues 306-319; and (K) residues 425-446 showing protective and deprotective effects of p52:p52 homodimer or (p52:p52):DNA on Bcl3-WT and Bcl3-4E. Bcl3-WT free protein was shown in grey; Bcl3-WT:(p52:p52) was shown in grey-dot; Bcl3-4E free protein was shown in blue; Bcl3-4E:(p52:p52) was shown in violet-dot; and Bcl3-4E:(p52:p52):DNA was shown in violet. All analyses were performed in experimental triplicate with error bar shown.

As mentioned above, there are no major changes in the deuterium uptake plots of Bcl3-4E in the Bcl3-4E:(p52:p52):DNA ternary complex compared to the Bcl3-4E:(p52:p52) binary complex barring the N-terminal 9-45 segment (Fig. 4D). Structural transition of this segment from rigid in free Bcl3-4E protein, to flexible in binary complex and back to rigid in the ternary complex is intriguing. The DNA-induced structural stability is allosteric in nature which might facilitate recruitment of selective factor(s) for transcription regulation. However, barring this segment, the difference uptake plot (Bcl3-4E:(p52:p52):DNA – Bcl3-4E:(p52:p52)), which accounts for the impact of DNA on the Bcl3-4E:(p52:p52) binary complex, showed small alterations throughout the rest of the protein. Moving along the Bcl3-4E molecule from residues 9-45 towards its C-terminus reveals that regions showing protection (residues 60-115 and 320-446) in the binary complex are deprotected in the ternary complex; while other region showing deprotection (residues 120-265) in binary complex is protected in the ternary complex; and the segment 265-320 showed less protection in the ternary complex compared to the binary complex (Fig. 4C-D). The deuterium uptake plots 5-minute timepoint of the last C-terminal segment of Bcl3-4E (residues 425-446) in its free form and as a ternary complex are comparable, whereas it is protected in the binary complex with p52 (Fig. 4K). Since the Bcl3 C-terminus positions close to the DNA binding region of p52 interfering with DNA binding, higher deuterium exchange of the C-terminus in the ternary complex suggests its exclusion from the DNA binding surface.

### Changes in dynamics of p52 within the (p52:p52):DNA and Bcl3:(p52:p52) binary complexes and Bcl3:(p52:p52):DNA ternary complex

To understand the altered dynamics of p52 as it transits from a free dimeric protein to a binary complex with Bcl3-4E and a ternary complex with Bcl3-4E and DNA together, we compared deuterium uptakes of p52 within these complexes. Bcl3 interacts asymmetrically with two identical protomers of p52 within its homodimer (Fig. 1E-F), similar to the binding mode observed for the interaction of IκBα with the p50:RelA heterodimer (Huxford et al. 1998; Ramsey et al. 2017). Thus, the protection or deprotection maps observed reflect an average of two specific interaction patterns. p52 binds DNA using five loops (L): L1 (45-78), L2 (140-145), L3 (220-229), L4 (251-258) and L5 (274-279) (PDB 1A3Q and 7CLI) (Cramer et al. 1997; Pan et al. 2023). The DNA binding protects p52 from deuterium uptake in three large regions: residues 30-90, 150-190 and 230-310 (Fig. 5A, 5E-F). In each of these regions, some segments were more protected than others, and some show no change. The DNA contacting residues in loops L2 and L3 appeared to be unprotected (Supplemental Fig. S3). Residues emanating from these two loops contact the center of DNA asymmetrically; loop L2 or L3 from one subunit of the dimer is stable and the other is flexible. In HDX-MS experiments of the (p50:RelA):DNA complex, these loops of the RelA subunit appeared unprotected, while the corresponding loops in p50 were protected (Narang et al. 2018). We suggest that the appearance of a lack for protection of these loops in the (p52:p52):DNA complex might be the result of asymmetric binding. Interestingly, there is only a limited number of peptides of p52 was protected upon DNA binding, and a few of these are the DNA binding loops L1, L4 and L5. A segment within GRR (residues 377-382), which is located away from the DNA binding surface, is deprotected upon DNA binding (Fig. 5G), similar to that observed in the case of p50 C-terminus in the (p50:RelA):DNA complex.

**Figure 5.**
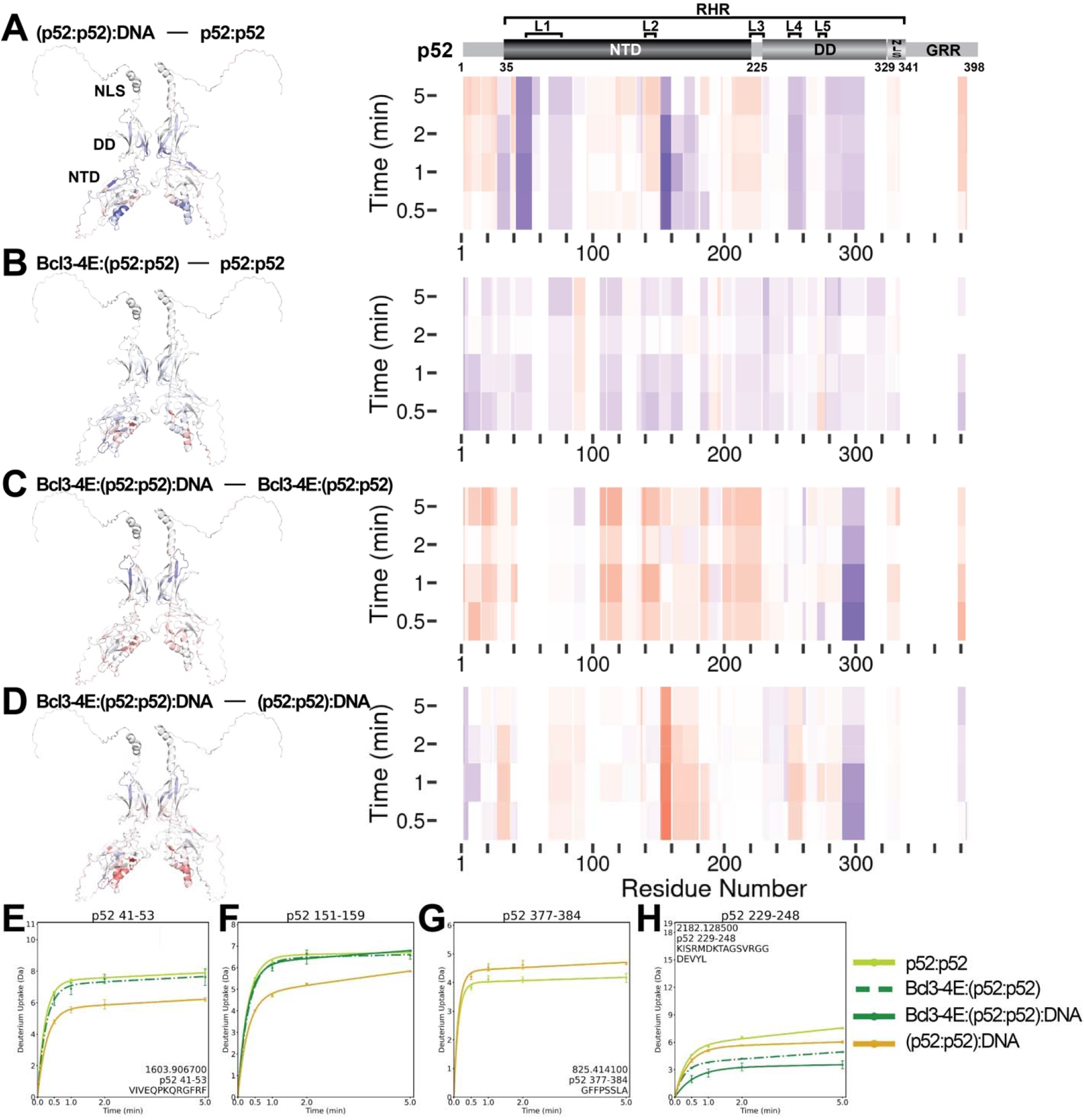
Dynamics changes of p52 homodimer upon binding to DNA and/or Bcl3-4E. The difference in fractional uptake of deuterium in p52:p52 homodimer (A) in DNA-bound vs. unbound states; (B) in Bcl3-4E-bound vs. unbound states; (C) in Bcl3-4E- and DNA-co-bound vs. Bcl3-4E-bound states; (D) in Bcl3-4E- and DNA-co-bound vs. DNA-bound states. The differences in fraction uptake displayed on p52 homodimer structural model are shown on the left; sequence ascribed heatmaps are shown on the right. Changes are represented in Blue-White-Red spectra (with grey representing absence of data, blue signifies protection, and red signifies deprotection), the indicated range of which is based on the spread of the differences (Supplemental Fig. S4). Uptake plot panels of representative p52 peptides (E) residues 41-53; and (F) residues 151-159 showing protective effect upon DNA binding. (G) Uptake plot panel of p52 peptide, residues 377-384, showing deprotective effect upon DNA binding. (H) Uptake plot panel of p52 peptide, residues 229-248, showing protective effect of upon both DNA and Bcl3-4E binding. The p52:p52 homodimer was shown in light green; Bcl3-4E:(p52:p52) was shown in dark green dash-dot; Bcl3-4E:(p52:p52):DNA was shown in dark green; and (p52:p52):DNA was shown in golden color. All analyses were performed in experimental triplicate with error bar shown.

Bcl3-4E protects the entire p52 in the Bcl3-4E:(p52:p52) binary complex, barring a few short segments (residues 90-120, 170-180, 190-200 and 275-280) (Fig. 5B). The protected regions include all the DNA binding loops L1-L5. The protection of these loops by Bcl3-4E, in particular, the region spanning residues 30-90, which contains the sequence specific DNA binding loop L1, poses a challenging concern as to how DNA binding is maintained in the Bcl3-4E:(p52:p52):DNA ternary complex. Only some regions protected by Bcl3-4E in the binary complexes remained to be protected in the ternary complex (Fig. 5B-C). And only two peptides, residues 229-248 (Fig. 5H) and residues 289-306 (Supplemental Fig. S3), show enhanced protection in the ternary complex as compared to that in the binary complexes. Of these, residues 229-248 locates right next to loop L3 which is engaged in DNA binding.

The differential protection map reveals that while the presence of Bcl3-4E in the ternary complex renders p52 globally more dynamic (Fig. 5D), this complex possesses certain features alluding to its unique ternary complex formation property. Indeed, the p52 protection map in the ternary complex showed that the DNA binding loops within its NTD were mostly exposed, suggesting that DNA binding might occur transiently. The greatest protection is observed in p52 DD, residues 289-306 (Supplemental Fig. S3). This segment was not protected by Bcl3-4E and was only partially protected by DNA in the binary complexes (Fig. 5A-B). Another segment in p52 NTD, spanning residues 85-95, showed deprotection in both binary complexes, and the deprotection was lost in the ternary complex (Fig. 5C-D). This result suggests that Bcl3 reorganizes p52’s interactions with DNA in its NTD, likely antagonizing binding of p52 to DNA. When p52 is bound to DNA, the effect of Bcl3-4E on p52 is largely similar with a couple of striking differences. Bcl3-4E enhanced deprotection of p52 NTD residues 152-168 and p52 DD residues 275-285 which contains the DNA-binding loop L5; however, the protection of residues 289-307 remained similar upon Bcl3-4E binding (Fig. 5D).

### The mechanism of phosphorylation-mediated transcriptional regulatory switch of Bcl3

The HDX data suggests that phosphorylation at S366 and S446 triggers altered conformation of the C-terminal flexible region of Bcl3 (Fig. 2), with apparent loss of several contacts with p52 NTD. It is possible that some of the loss in interactions is partly compensated by new contacts, possibly between Bcl3 ARD and the βe-βf loop (289-306) within p52 DD (Fig. 5C-D; Supplemental Fig. S3). The βe-βf loop residues are distinct among members of the NF-κB family (Fig. 6A), hinting such contacts to be specific for NF-κB p52. It has been reported that p52 interacts with another transcription factor ETS1 via residues in this loop to activate the telomerase reverse transcriptase (*TERT*) promoter (Xu et al. 2018). The structural model of the Bcl3:(p52:p52):DNA complex presented in Fig. 1E and 1F indicates that the βe-βf loop is located near Bcl3 AR3, suggesting the possibility of contacts between the two proteins via this loop. To test this, we replaced the βe-βf loop sequence of p52 with its counterparts from p50 (named p52-KDIN) and RelA (named p52-ADPS), and performed a pulldown experiment using biotinylated P-Selectin-κB DNA with equal amounts of p52-WT or mutant p52 (p52-ADPS and p52-KDIN) alone, Bcl3-4E alone, or a mixture with p52-WT, p52-ADPS or p52-KDIN mutants (Fig. 6B). All the p52 proteins were purified to similar purity (Supplemental Fig. S6A). We found that the p52-KDIN mutant retained lesser amounts of Bcl3-4E compared to p52-WT. However, the p52-ADPS mutant showed no significant difference. Although this p52-ADPS mutant appeared to bind the DNA better than p52-WT in the absence of Bcl3-4E. We further tested if these mutants were less effective in driving transcription. We observed significantly reduced transcriptional activity of both p52 mutants in a reporter-based assay (Fig. 6C). These results suggest that the βe-βf loop of p52 plays a role in transcription, likely through its involvement in Bcl3 binding to drive transcription. These results also hint at the possibility of Bcl3 acting as an allosteric regulator of p52, dictating the nature of the molecular interaction between p52 and DNA.

**Figure 6.**
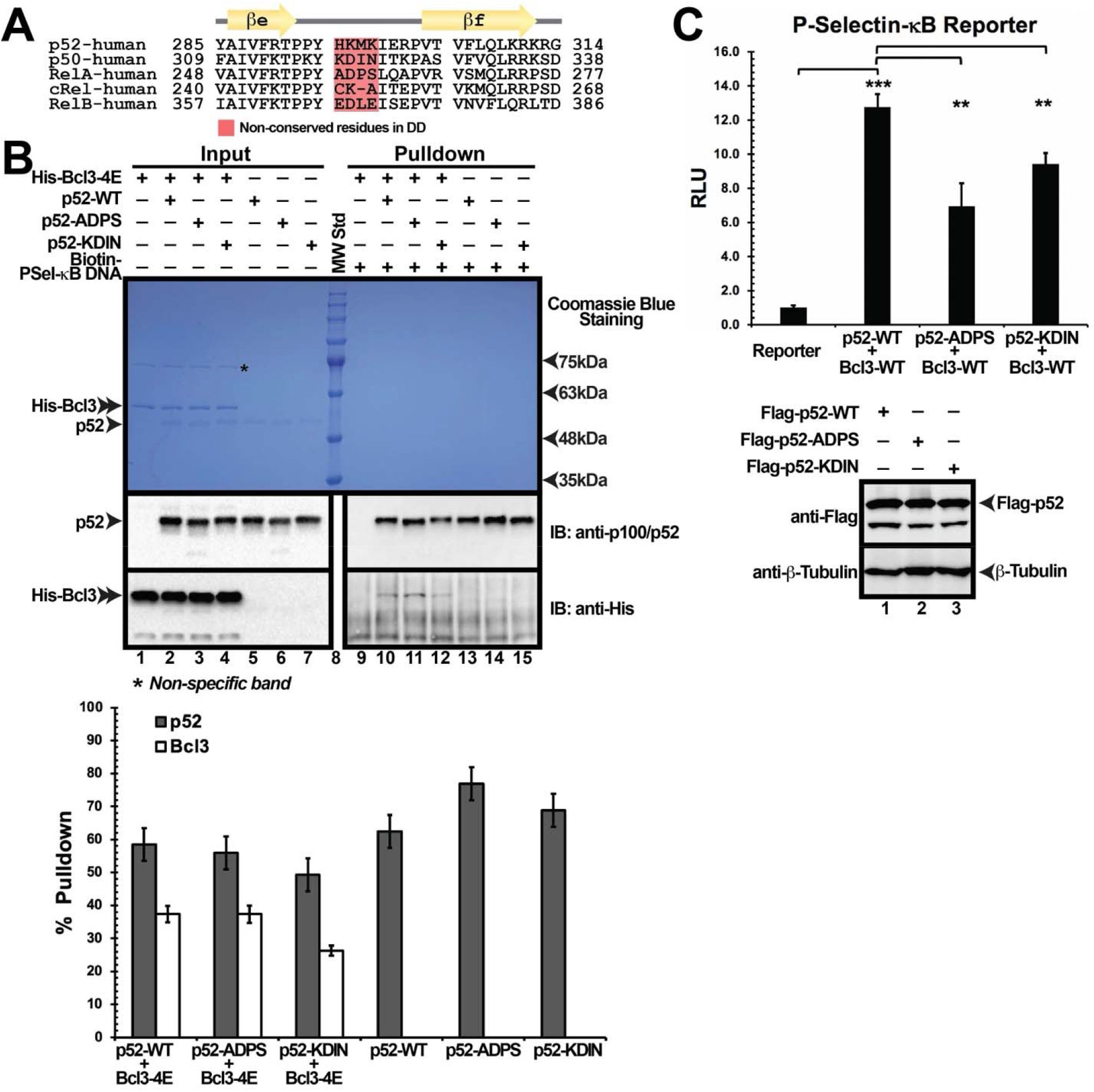
p52 βe-βf loop plays a role in transcriptional regulation. (A) Sequence alignment of the βe-βf loop of NF-κB subunits. Secondary structures and connecting loops are drawn above the sequences. (B) Top: Streptavidin pulldown assay demonstrating the binding of recombinant p52-WT and -ADPS, -KDIN mutant proteins alone or together with His-Bcl3-4E to the biotinylated P-Selectin-κB DNA. Bottom: Plots showing the average amount of indicated protein bound to DNA from three independent experiments at the same condition, error bar represent SD. (C) Top: Luciferase reporter activity driven by co-expression of Bcl3 and p52 and its hybrid mutants indicates significant reduction in transcriptional activity with mutation. The data were analyzed from three independent experiments performed in triplicate. RLU, relative luciferase unit. **p<0.01; ***p<0.001 (t test). Error bars represent SD. Bottom: Western blot assay showing similar level of expression of p52 or its mutants in cells.

Furthermore, the changes in deuterium uptake properties of both Bcl3 and p52 strongly suggest that the C-terminal flexible region of Bcl3 is juxtaposed with the p52 NTD and makes direct contacts, which results in the inhibition of DNA binding by p52. Phosphorylation of Bcl3 at S366 and S446 alleviates this inhibitory effect on DNA binding by p52. This prompted us to test if the deletion of Bcl3 C-terminal segment could relieve its inhibitory activity towards forming a ternary complex. We generated a truncation mutant of Bcl3 lacking 28 C-terminal residues (Bcl3-WT^1-418^) and performed the pulldown experiment using biotinylated P-Selectin-κB DNA with equal amounts of phospho-mimetic Bcl3-4E, unphosphorylated Bcl3-WT or the C-terminal truncation mutant Bcl3-WT^1-418^ alone, p52-WT alone, or a mixture with Bcl3-4E, Bcl3-WT or Bcl3-WT^1-418^ mutants. All the Bcl3 proteins were expressed and purified to similar purity (Supplemental Fig. S6B). The result shows that the unphosphorylated full-length Bcl3-WT formed ternary complex with p52 and DNA weakly; however, the unphosphorylated Bcl3-WT^1-418^ was able to form the ternary complex comparable to the phospho-mimetic Bcl3-4E (Fig. 7A). Furthermore, the luciferase reporter assay with co-expression of Bcl3-WT^1-418^ together with p52 showed higher luciferase activity compared to Bcl3-WT in cells (Fig. 7B). It should be noted that unlike the unphosphorylated recombinant Bcl3-WT protein expressed in *E. coli*, Bcl3-WT is constitutively phosphorylated when expressed in mammalian cells and is able to induce expression of reporter genes. However, extent of phosphorylation at S366 and S446 could vary, leading to differential outcomes. The removal of Bcl3 C-terminal tail abrogated its inhibitory effect, irrespective of the phosphorylation state at S366. These finding support our hypothesis that the C-terminal region of Bcl3 is inhibitory to the ternary complex formation. Interestingly, most of the oncogenic mutations are located within this flexible C-terminus starting from residue 394 emphasizing the functional significance of this segment (Figure 7C). Phosphorylation at both S366 and S446 likely renders structural alteration and increased flexibility of Bcl3 C-terminus, thereby allowing the access of p52 to target DNA while p52 remains bound to Bcl3 as depicted in a model shown in Fig. 7D. Collectively, these results suggest tight regulation of phosphorylation is important for Bcl3’s normal function as a transcriptional coactivator and its aberration due to mutations lead to oncogenic activity.

**Figure 7.**
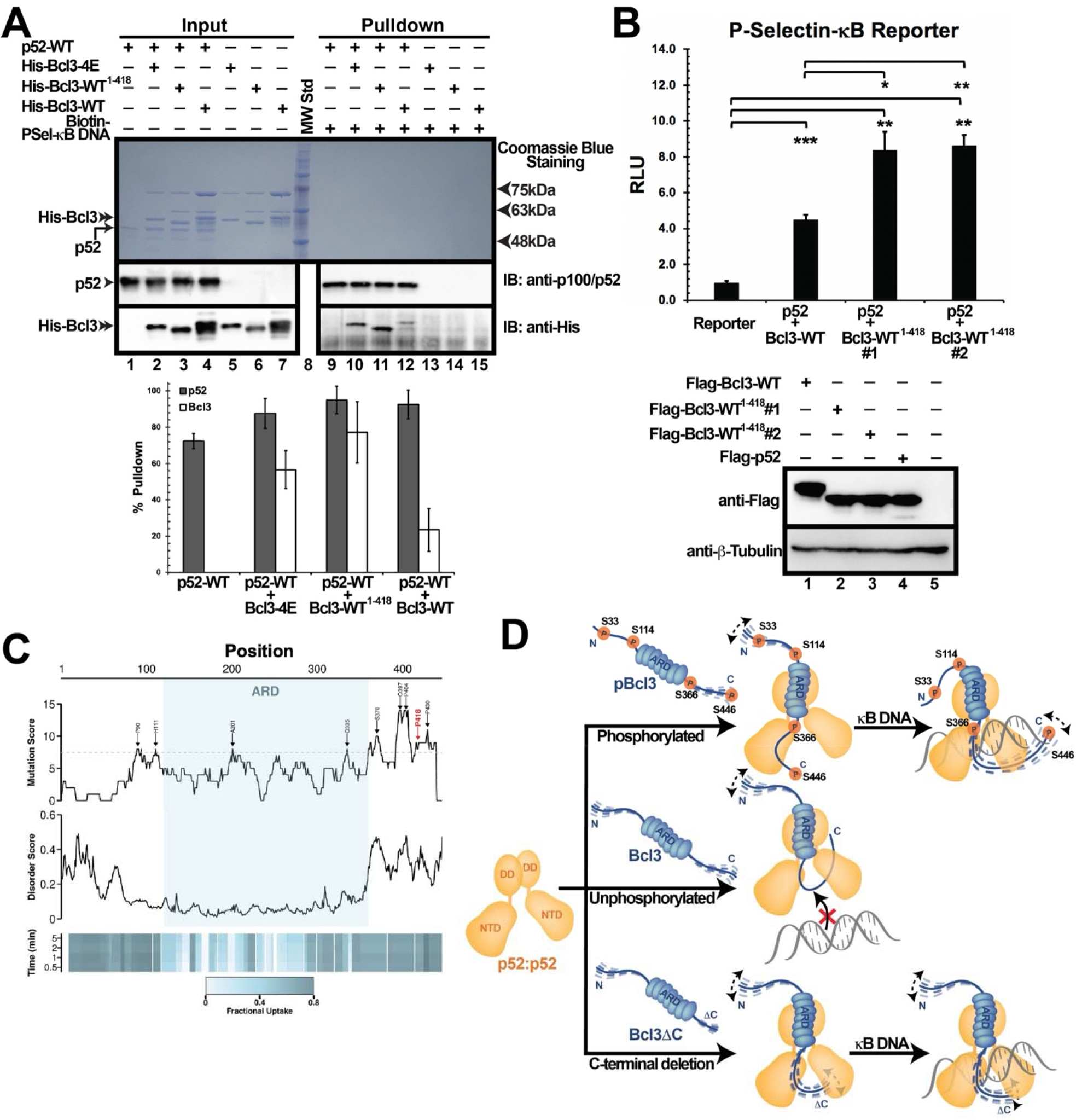
Phosphorylation-mediated transcriptional regulatory switch of Bcl3. (A) Top: Streptavidin pulldown assay demonstrating the binding of recombinant His-Bcl3-4E, His-Bcl3-WT, and His-Bcl3-WT^1-418^ proteins alone or together with p52-WT to the biotinylated P-Selectin-κB DNA. Bottom: Plots showing the average amount of indicated protein bound to DNA from three independent experiments at the same condition, error bar represent SD. (B) Top: Luciferase reporter activity driven by co-expression of p52 and Bcl3-WT or Bcl3-WT^1-418^ indicating enhanced transcriptional activation C-terminal truncation of Bcl3. Two equivalent clones of Bcl3-WT^1-418^ were tested. The data were analyzed from three independent experiments performed in triplicate. RLU, relative luciferase unit. *p<0.05; **p<0.01; ***p<0.001 (t test). Error bars represent SD. Bottom: Western blot assay showing similar level of expression of Bcl3-WT and uponz Bcl3-WT^1-418^ in cells. (C) Top: Plot representing substitution mutation frequency of Bcl3 identified in cancers from COSMIC database. A sliding window of 15 amino acids was used for counting mutations across Bcl3. Dotted line represents an approximate midpoint mutation rate, and labeled residues denote regions of elevated mutation frequency. P418 is shown in red. Middle: Plot of predicted disorder of Bcl3 using flDPnn tool. Bottom: Heatmap representation of fractional deuterium uptake for Bcl3-WT at indicated timepoints. (D) A cartoon model showing the transcriptional regulation by Bcl3 phosphorylation via NF-κB p52:p52 homodimer DNA binding.

## Discussion

This study explores how Bcl3, a member of the IκB family proteins, functions as a regulator of p52:p52 homodimer mediated transcriptional activation. It has long been known that Bcl3 requires phosphorylation to function as a transcriptional coactivator (Bundy and McKeithan 1997). In this study, we show that phosphorylation at four residues, two located at the N-terminal (S33, S114) and other two at the C-terminal flexible region (S366, S446) of Bcl3, regulates the formation of a transcriptionally competent Bcl3:(p52:p52):DNA ternary complex. Phosphorylation at S33 is essential for both stabilization and nuclear localization of Bcl3; however, it is unclear if phosphorylation at any of the other three sites is strictly required for a specific or all transcription events mediated by the Bcl3:(p52:p52) complex. Several other residues of Bcl3 also undergo phosphorylation although their roles in transcription have not been thoroughly investigated. It is possible that phosphorylation of a specific set of residues induces transcription of a specific gene.

We also observed that the C-terminal 28-residue segment of Bcl3 is inhibitory to the formation of the ternary complex. Phosphorylation at both S366 and S446 allow positioning of this segment away from the p52 and DNA binding interface. The S366 has been observed to be mutated in various cancers and inflammatory diseases (COSMIC database legacy identifier number COSM9762159 and COSM10046017) (Bonilla et al. 2016; Yuan et al. 2016). This reflects significance of S366 in regulating activity of Bcl3, and its phosphorylation linked to pathogenicity. Further studies are required to test if phosphorylation of different sets of residues within the C-terminal flexible region demarcates transcriptional activity of Bcl3 under pathogenic vs. non-pathogenic conditions.

Our HDX-MS results revealed that phosphorylation of Bcl3 renders it more dynamic, with enhanced deuterium uptake throughout the molecule except for its N-terminal 9-45 region. This dynamicity persists even upon its binding to the p52:p52 homodimer or the (p52:p52):DNA complex. Only a few additional contacts between p52 and DNA are observed when Bcl3-4E engages to the (p52:p52):DNA complex. Rather, a number of specific and high-energy contacts between p52 and DNA are lost, suggesting that Bcl3-4E weakens the interaction between p52 and DNA. This counterintuitive results hints at more complex possibilities of Bcl3 mediated transcriptional activation such as assembly of specific protein factors with this ternary complex leading to a stabilized multiprotein transcriptional complex. However, we cannot rule out the possibility that the transcriptional complexes need to be weak and unstable for gene synthesis *in vivo*. Consistent with our observations, other IκB proteins shown to be involved in transcription activation in cells by forming ternary complexes with NF-κB and DNA also dislodge DNA from NF-κB *in vitro*. A recent publication of IκBζ in complex with p50:p50 homodimer reveals AR7 of IκBζ clashes severely with DNA and it is proposed that the extreme C-terminus of IκBζ mediates structural rearrangement to accommodate DNA (Zhu et al. 2024). Future experiments are required to relate the stability of transcription factor - DNA complexes to gene synthesis.

Our study also reveals dynamicity of Bcl3 N-terminal 9-45 region - altering from rigid in its free form, to flexible in (p52:p52)-bound state, and back to rigid in the (p52:p52):DNA-bound state. The reduced deuterium exchange of this region within the ternary complex suggests that cognate DNA possibly induces allosteric conformational changes. This region is reported to recruit ubiquitin ligase, TBL/TBLR1, in the cytoplasm when S33 is unphosphorylated to induce Bcl3 degradation (Keutgens et al. 2010; Wang et al. 2017). There are reports that a class of Ub ligase interacts with HDAC3 and that an overlapping region also recruits transcriptional corepressor CtBP (Keutgens et al. 2010; Milazzo et al. 2020). These latter events occur in the nucleus targeting S33 phosphorylated Bcl3. However, the cellular conditions for binding these repressors have not been investigated. In this study, we used the natural G/C-centric P-Selectin-κB DNA, which is known to recruit Tip60, a histone acetylases, to the Bcl3:(p52:p52) complex and activate transcription (Wang et al. 2012). It is possible that an A/T-centric κB DNA would alter the conformation of this region differently when it binds to the Bcl3:(p52:p52) binary complex, recruiting histone deacetylase and repressing transcription. Collectively, these observations led us to speculate that distinct conformations of the N-terminal region of Bcl3 upon its phosphorylation and/or binding to specific DNA sequences regulate the selection of protein factor such as TBLR1, CtBP, HDAC, and possibly Tip60 conferring specific activities.

Bcl3 is an oncoprotein, which has been implicated in a wide range of cancers, both hematological and solid tumors. The constitutive nuclear expression of Bcl3 induces chronic inflammation and proliferation by maintaining high levels of cyclin D1 and inflammatory cytokines, controlling both tumor growth and proliferation. In most cases where an aberrant Bcl3 is responsible for the cancer phenotype, it is overexpressed. Bcl3 is also mutated in many cases. The effects of these mutations are yet to be fully understood, although a significant number of these mutations are clustered in its C-terminal regions (COSMIC database). Our results indicate that inappropriate phosphorylation or a C-terminal deletion of Bcl3 could affect expression of its regulatory genes mediating inflammation and proliferation.

## Materials and Methods

### Protein Expression and Purification

Recombinant non-tagged human p52 (1-398)-WT, -ADPS and -KDIN mutants were expressed and purified from *Escherichia coli* Rosetta (DE3) cells. Rosetta (DE3) cells transformed with pET-11a-p52 (1-398)-WT, -ADPS or -KDIN mutants were cultured in 2 L of LB medium containing 50 mg/mL ampicillin and 34 mg/mL chloramphenicol at 37 ^o^C. Expression was induced with 0.2mM Isopropyl β-D-1-thiogalactopyranoside (IPTG) at OD_600_ 0.5-0.6 for 3 hours. Cells were harvested by centrifugation, suspended in 40 mM Tris-HCl pH 7.5, 100 mM NaCl, 10 mM β-Mercaptoethanol (β-ME), 1 mM PMSF, and lysed by sonication. Cell debris was removed by centrifugation (20,000 g for 30 min). Clarified supernatant was loaded onto Q-Sepharose FF column (GE Healthcare). Flow-through fraction was applied to SP HP column (GE Healthcare). The column was washed with 40 mM Tris-HCl pH 7.5, 200 mM NaCl; 10 mM β-ME, and the protein was eluted by the same buffer containing 400 mM NaCl. p52 was concentrated and loaded onto the gel filtration column (HiLoad 16/600 Superdex 200 pg, GE Healthcare) pre-equilibrated with 10 mM Tris-HCl (pH 7.5), 100 mM NaCl; 5 mM β-ME. Peak fractions were concentrated to ∼10 mg/mL, flash frozen in liquid nitrogen and stored at -80°C.

Recombinant human His-Bcl3 (1-446)-WT, phospho-mimetic mutants, and (1-418) C-terminal deletion mutant were expressed in *Escherichia coli* Rosetta (DE3) cells by induction with 0.2 mM IPTG at OD_600_ 0.4 for 8 hours at 24°C. Cell pellets of 2 L-culture of Bcl3 alone or together with 1 L-culture of p52 (for p52:p52:Bcl3 complex) were resuspended together in buffer containing 20 mM Tris-HCl pH 8.0, 300 mM NaCl, 25 mM imidazole, 10% (v/v) glycerol, 10 mM β-ME, 0.1 mM PMSF and 50μL protease inhibitor cocktail (Sigma, Cat#P8465) and then purified by Ni Sepharose (HisTrap HP, GE) followed by anion exchange column (Q Sepharose fast flow, GE). The protein complexes further went through HiTrap Desalting Column (GE) to exchange buffer before other assays.

### Luciferase Reporter Assays

HeLa cells were obtained from ATCC (CCL-2) and cultured in Dulbecco’s modified Eagle’s medium (DMEM; Gibco, Cat#11995065) that was supplemented with 10% fetal bovine serum (FBS; Gibco, Cat#10270106) and 1x Penicillin-Streptomycin-L-Glutamine (Corning, Cat#30-009-Cl). HeLa cells were transiently transfected with Flag-p52(1-415)-WT (or various mutants) together with Flag-Bcl3(1-446) (or various mutants) expression vectors or empty Flag-vector, and the luciferase reporter DNA with P-Sel-κB DNA promoter (Wang et al. 2012). The total amount of plasmid DNA was kept constant for all assays. Transient transfections were carried out using polyethylenimine (PEI) (Polysciences, Cat#23966). Cells were collected 48 hours after transfection. Luciferase activity assays were performed using Dual-Luciferase Reporter Assay System (Promega, Cat#E1910) following the manufacturer’s protocol. Data are represented as mean standard deviations (SD) of three independent experiments in triplicates.

### Electrophoretic Mobility Shift Assays (EMSA)

The oligonucleotides containing κB site (Wang et al., 2012) used for EMSA were PAGE purified, end radiolabeled with ^32^P using T4-polynucleotide kinase (NEB, Cat#M0201) and [γ-^32^P] ATP, and annealed to the complementary strands. Binding reaction mixtures contained indicated recombinant proteins or nuclear extracts at different concentrations, binding buffer (10mM Tris-HCl pH 7.5, 50mM NaCl, 10% (v/v) glycerol, 1% (v/v) NP-40, 1mM EDTA, 0.1mg/mL poly(dI-dC)), and ∼10000cpm of ^32^P-labeled DNA. Reaction mixtures were incubated at room temperature for 20 minutes and analyzed by electrophoresis on a 5% (w/v) non-denatured polyacrylamide gel at 200V for 1 hour in 25mM Tris, 190mM glycine, and 1mM EDTA. The gels were then dried, exposed to a phosphorimager, and scanned by Typhoon scanner 9400 (Amershan Bioscience). Gels were quantified by using ImageQuant version5.2 (Molecular Dynamics).

### Streptavidin Pulldown Assays

100 nM annealed biotinylated P-Selectin-κB DNA was pre-incubated with 15 μL streptavidin magnetic beads (MedChemExpress, Cat#HY-K0208) using incubation buffer containing 10 mM Tris pH 7.5, 1 mM EDTA, 1 M NaCl, 0.02% (v/v) Tween-20 at 4°C for 1 hour followed by washing with 1 mL incubation buffer. 80 ng of recombinant non-tagged p52(1-398)-WT or -ADPS, -KDIN mutant protein were mixed with recombinant 0.8 μg His-Bcl3(1-446)-4E or His-Bcl3(1-446)-WT, His-Bcl3(1-418) protein in binding buffer containing 20 mM Tris pH 7.5, 100 mM NaCl, 0.1% (v/v) NP-40, and 1 mM DTT. Complexes of p52 with His-Bcl3 were then pull-downed using 15 μL Biotin-κB DNA pre-incubated streptavidin magnetic beads at 4°C for 2 hours. Bound complexes were washed three times with binding buffer and dissolved in 4xSDS-dye by boiling at 95°C for 10 minutes followed by SDS-PAGE. Bound p52 and His-Bcl3 proteins were visualized by immunoblotting using anti-NF-κB p100/p52 antibody (Cell Signaling Technology, Cat#4882) and anti-His antibody (Bio Bharati Life Science, Cat#BB-AB0010) respectively. The pulldown assays were performed in experimental triplicate, immunoblotting images were then analyzed using Image-J software to measure the band intensity of p52 and Bcl3 proteins.

### Hydrogen-Deuterium Exchange Mass Spectrometry (HDX-MS)

Bcl3-WT or Bcl3-4E and p52 (i.e., p52:p52 homodimer) were exchanged to HDX buffer (300 mM NaCl, 25 mM Tris pH 8.0, 5 % glycerol, 10 mM β-mercaptoethanol) prior to HDX-MS experiments. Commonly, a 100 μL volume of concentrated protein was loaded onto a Superose6 5/150 increase column (Cytiva), and appropriate elution peak fractions from multiple runs were combined and concentrated using a protein concentrator (Sartorius, Vivaspin, 30,000 MWCO) to obtain a final sample of ∼300 μL at ∼30 μM concentration.

For HDX-MS of Bcl3 and p52 homodimer, proteins at 10 μM concentration in HDX buffer were used. When HDX-MS of Bcl3 and p52 were performed in the presence of Ub_4_, a three-fold molar excess of Ub_4_ (i.e., ∼30 μM) was present.

For HDX-MS of Bcl3:(p52:p52) complex, an equimolar mixture of Bcl3 and p52 homodimer was HDX-buffer exchanged using a Superose 6 5/150 increase column. Peak fractions of the Bcl3:(p52:p52) complex from multiple runs were pooled and concentrated, and the concentration was adjusted to ∼5 μM. When HDX-MS assays of Bcl3:(p52:p52) was performed in presence of Ub_4_, the concentration of Bcl3:(p52:p52) complex was ∼3 μM and that of Ub_4_ was ∼9 μM.

HDX-MS was performed at the Biomolecular and Proteomics Mass Spectrometry Facility (BPMSF) of the University California San Diego, using a Waters Synapt G2Si system with HDX technology (Waters Corporation) as described previously (Ramirez-Sarmiento and Komives 2018; Vinciauskaite and Masson 2022). Deuterium exchange reactions were performed using a Leap HDX PAL autosampler (Leap Technologies, Carrboro, NC). The HDX-buffer for deuteration contained 300 mM NaCl, 25 mM Tris pH 8.0, 5% glycerol, and 10 mM β-mercaptoethanol in D_2_O. D_2_O was exchanged for H_2_O by lyophilizing buffers initially prepared in ultrapure water and redissolving in the same volume of 99.96% D_2_O (Cambridge Isotope Laboratories, Inc., Andover, MA) immediately before use. Measurements were made in triplicate for each deuteration timepoint analyzed (0.5, 1, 2, and 5 min). For each reaction, a 4 μL volume of sample (at the abovementioned concentrations) was stabilized at 25°C for 5 min prior to mixing with 56 μL of D_2_O containing HDX-buffer. The reaction was quenched for 1 min at 1°C by combining 50 μL of the reaction with 50 μL of 3 M guanidine hydrochloride, final pH 2.66. The quenched sample (90 μL) was then injected in a 100 μL sample loop, followed by digestion on an in-line pepsin column (Immobilized Pepsin, Pierce) at 15°C. The resulting peptides were captured on a BEH C18 Vanguard precolumn, separated by analytical chromatography (Acquity UPLC BEH C18, 1.7 μm 1.0 × 50 mm, Waters Corporation) using a 7–85% acetonitrile gradient in 0.1% formic acid over 7.5 min, and electrosprayed into the Waters Synapt G2Si quadrupole time-of-flight mass spectrometer. The mass spectrometer was set to collect data in the Mobility, ESI+ mode; mass acquisition range of 200–2000 (m/z); scan time 0.4 s. Continuous lock mass correction was accomplished with infusion of leu-enkephalin (m/z = 556.277) every 30 s (mass accuracy of 1 ppm for calibration standard).

For peptide identification, the mass spectrometer was set to collect data in mobility-enhanced data-independent acquisition (MS^E^), mobility ESI+ mode. Peptide masses were identified from triplicate analyses and data were analyzed using the ProteinLynx global server (PLGS) version 3.0 (Waters Corporation). Peptide masses were identified using a minimum number of 250 ion counts for low energy peptides and 50 ion counts for their fragment ions; the peptides also had to be larger than 1,500 Da. The following cutoffs were used to filter peptide sequence matches: minimum products per amino acid of 0.2, minimum score of 7, maximum MH+ error of 5 ppm, and a retention time RSD of 5%. In addition, the peptides had to be present in two of the three ID runs collected. The peptides identified in PLGS were then analyzed using DynamX 3.0 data analysis software (Waters Corporation). The relative deuterium uptake for each peptide was calculated by comparing the centroids of the mass envelopes of the deuterated samples with the undeuterated controls following previously published methods(Ramirez-Sarmiento and Komives 2018; Lumpkin and Komives 2019). For all HDX-MS data, 1 biological replicate was analyzed with 3 technical replicates. Data represented from different biological replicates are noted in their corresponding figures. All data are represented as mean values +/-SEM of 3 technical replicates due to limitations of processing software, however the LEAP robot provides highly reproducible data for biological replicates. The deuterium uptake was corrected for back-exchange using a global back exchange correction factor determined from the average percent exchange measured in disordered regions of each protein (Ramsey et al. 2017).

### HDX-MS Data Analysis

The DynamX 3.0 (Waters) output of HDX-MS data for p52 and Bcl3 peptides (consolidated data in .csv format are provided as Supporting files) were processed using DECA version 1.16 (Ramirez-Sarmiento and Komives 2018; Lumpkin and Komives 2019) to calculate the fractional exchange compared to theoretical maximum at the residue level of individual peptides from individual states. The differences in fractional exchange of residues between pair of individual states were computed and plotted as a sequence ascribed heatmap using ‘R’. The ranges of the ‘difference in fractional exchange’ values were determined, and differences were represented as heatmaps of the blue-white-red color spectra to visualize the protection/deprotection. These ‘difference in fractional exchange’ values were also used to generate colored structural cartoons using the PyMOL Molecular Graphics System, Version 3.0 Schrödinger, LLC.

## Acknowledgments

This work was supported by the Science and Technology Development Fund, Macao S.A.R. (FDCT) [project 0104/2019/A2 and 0089/2022/AFJ to V.Y.-F.W.]; the Multi-Year Research Grant from University of Macau [MYRG2018-00093-FHS to V.Y.-F.W.]; and the National Institutes of Health (NIH) [GM085490 to G.G.].

## Author Contributions

V.Y.-F.W. and G.G. conceptualized and supervised the project. V.Y.-F.W. performed the HDX-MS experiments; T.B., S.S., and W.S. analyzed the HDX-MS data and generated the related figures. W.P. performed all the biochemistry and molecular biology experiments. Q.D. and A.V. performed the EMSA. V.Y.-F.W., G.G. and T.B. wrote the paper with input from the other authors.

