## Supplemental Information for "Phosphorylation-induced flexibility of proto-oncogenic Bcl3 regulates transcriptional activation by NF-κB p52 homodimer"

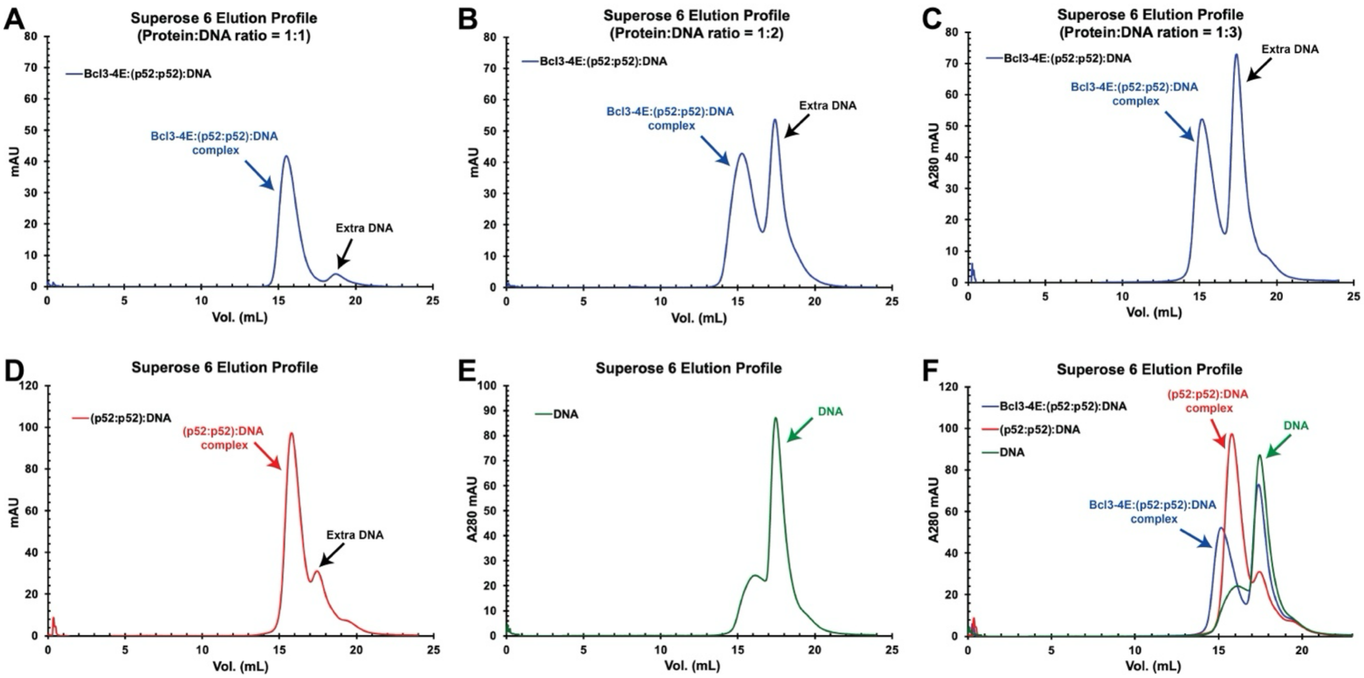


**Figure S1. Superose 6 size exclusion chromatography elution profile showing the ternary complex formation of Bcl3-4E:(p52:p52):DNA.** Elution profile of the ternary complex using protein:DNA ratio of (A) 1:1; (B) 1:2; (C) 1:3. Elution protein of (D) (p52:p52):DNA binary complex and (E) DNA alone. (F) Overlay of elution profiles of ternary complex, binary complex and DNA.

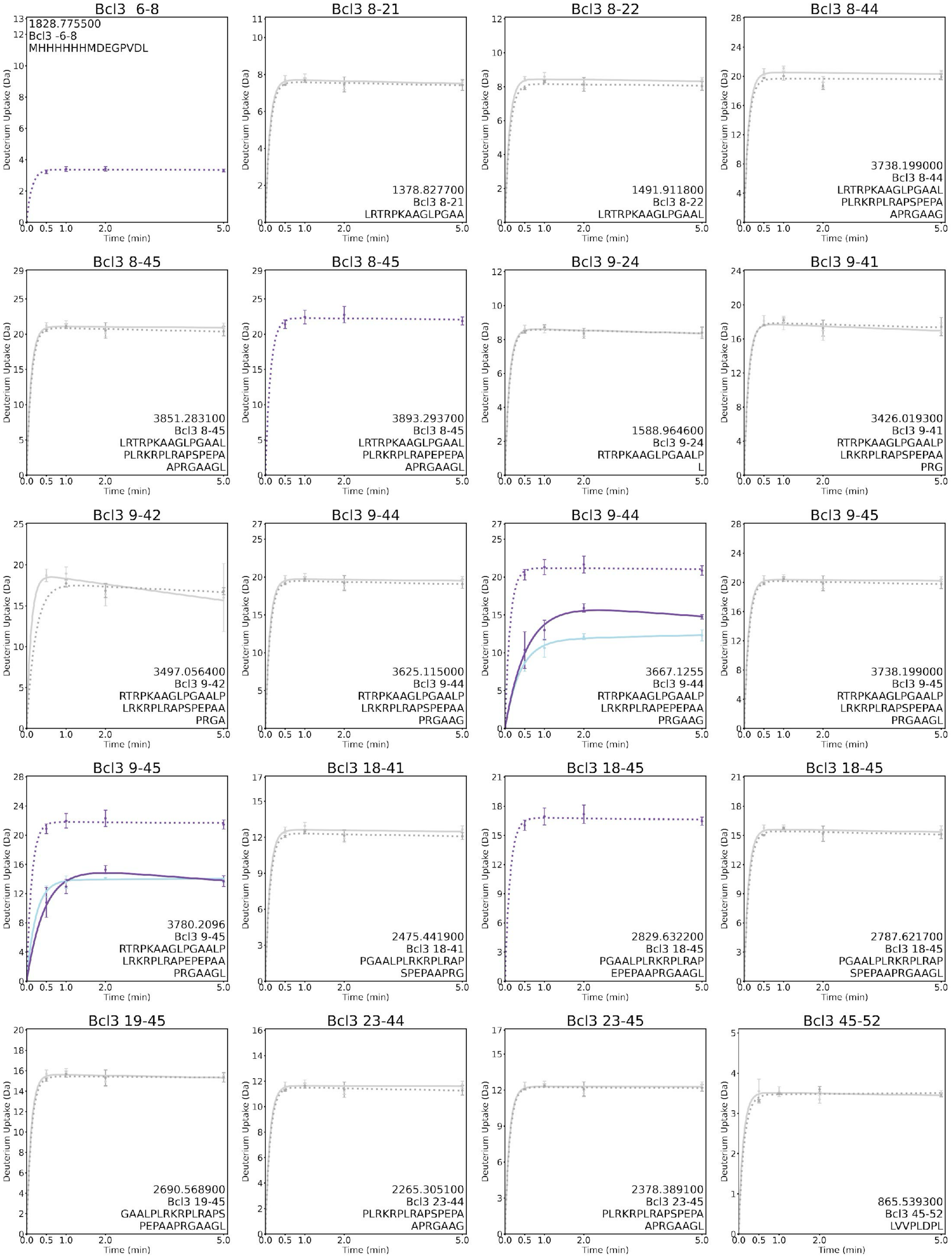

Bcl3 6-8
1828.775500
Bcl3 -6-8
MHHHHHHMDEGPVDL
Bcl3 8-21
1378.827700
Bcl3 8-21
LRTRPKAAGLPGAA
Bcl3 8-22
1491.911800
Bcl3 8-22
LRTRPKAAGLPGAAL
Bcl3 8-44
3738.199000
Bcl3 8-44
LRTRPKAAGLPGAAL
PLRKRPLRAPSPEPA
APRGAAG
Bcl3 8-45
3851.283100
Bcl3 8-45
LRTRPKAAGLPGAAL
PLRKRPLRAPSPEPA
APRGAAGL
Bcl3 8-45
3893.293700
Bcl3 8-45
LRTRPKAAGLPGAAL
PLRKRPLRAPEPEPA
APRGAAGL
Bcl3 9-24
1588.964600
Bcl3 9-24
RTRPKAAGLPGAALP
L
Bcl3 9-41
3426.019300
Bcl3 9-41
RTRPKAAGLPGAALP
LRKRPLRAPSPEPAA
PRG
Bcl3 9-42
3497.056400
Bcl3 9-42
RTRPKAAGLPGAALP
LRKRPLRAPSPEPAA
PRGA
Bcl3 9-44
3625.115000
Bcl3 9-44
RTRPKAAGLPGAALP
LRKRPLRAPSPEPAA
PRGAAG
Bcl3 9-44
3667.1255
Bcl3 9-44
RTRPKAAGLPGAALP
LRKRPLRAPEPEPAA
PRGAAG
Bcl3 9-45
3738.199000
Bcl3 9-45
RTRPKAAGLPGAALP
LRKRPLRAPSPEPAA
PRGAAGL
Bcl3 9-45
3780.2096
Bcl3 9-45
RTRPKAAGLPGAALP
LRKRPLRAPEPEPAA
PRGAAGL
Bcl3 18-41
2475.441900
Bcl3 18-41
PGAALPLRKRPLRAP
SPEPAAPRG
Bcl3 18-45
2829.632200
Bcl3 18-45
PGAALPLRKRPLRAP
EPEPAAPRGAAGL
Bcl3 18-45
2787.621700
Bcl3 18-45
PGAALPLRKRPLRAP
SPEPAAPRGAAGL
Bcl3 19-45
2690.568900
Bcl3 19-45
GAALPLRKRPLRAPS
PEPAAPRGAAGL
Bcl3 23-44
2265.305100
Bcl3 23-44
PLRKRPLRAPSPEPA
APRGAAG
Bcl3 23-45
2378.389100
Bcl3 23-45
PLRKRPLRAPSPEPA
APRGAAGL
Bcl3 45-52
865.539300
Bcl3 45-52
LVVPLDPL
Deuterium Uptake (Da)
Time (min)

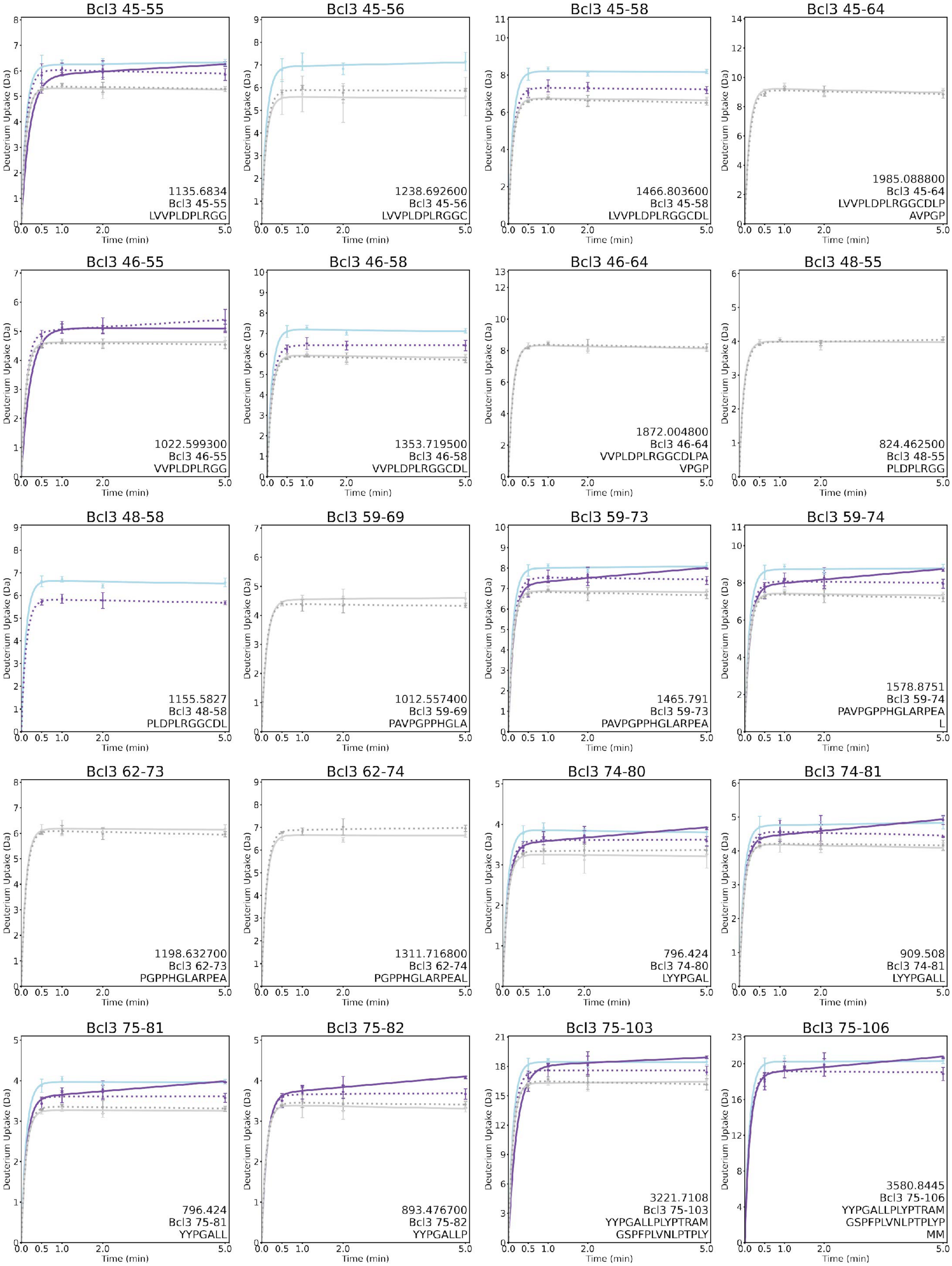
Bcl3 45-55
1135.6834
Bcl3 45-55
LVVPLDPLRGG
Bcl3 45-56
1238.692600
Bcl3 45-56
LVVPLDPLRGGC
Bcl3 45-58
1466.803600
Bcl3 45-58
LVVPLDPLRGGCDL
Bcl3 45-64
1985.088800
Bcl3 45-64
LVVPLDPLRGGCDLP
AVPGP
Bcl3 46-55
1022.599300
Bcl3 46-55
VVPLDPLRGG
Bcl3 46-58
1353.719500
Bcl3 46-58
VVPLDPLRGGCDL
Bcl3 46-64
1872.004800
Bcl3 46-64
VVPLDPLRGGCDLPA
VPGP
Bcl3 48-55
824.462500
Bcl3 48-55
PLDPLRGG
Bcl3 48-58
1155.5827
Bcl3 48-58
PLDPLRGGCDL
Bcl3 59-69
1012.557400
Bcl3 59-69
PAVPGPPHGLA
Bcl3 59-73
1465.791
Bcl3 59-73
PAVPGPPHGLARPEA
Bcl3 59-74
1578.8751
Bcl3 59-74
PAVPGPPHGLARPEA
L
Bcl3 62-73
1198.632700
Bcl3 62-73
PGPPHGLARPEA
Bcl3 62-74
1311.716800
Bcl3 62-74
PGPPHGLARPEAL
Bcl3 74-80
796.424
Bcl3 74-80
LYYPGAL
Bcl3 74-81
909.508
Bcl3 74-81
LYYPGALL
Bcl3 75-81
796.424
Bcl3 75-81
YYPGALL
Bcl3 75-82
893.476700
Bcl3 75-82
YYPGALLP
Bcl3 75-103
3221.7108
Bcl3 75-103
YYPGALLPLYPTRAM
GSPFPLVNLPTPLY
Bcl3 75-106
3580.8445
Bcl3 75-106
YYPGALLPLYPTRAM
GSPFPLVNLPTPLYP
MM
Deuterium Uptake (Da)
Time (min)

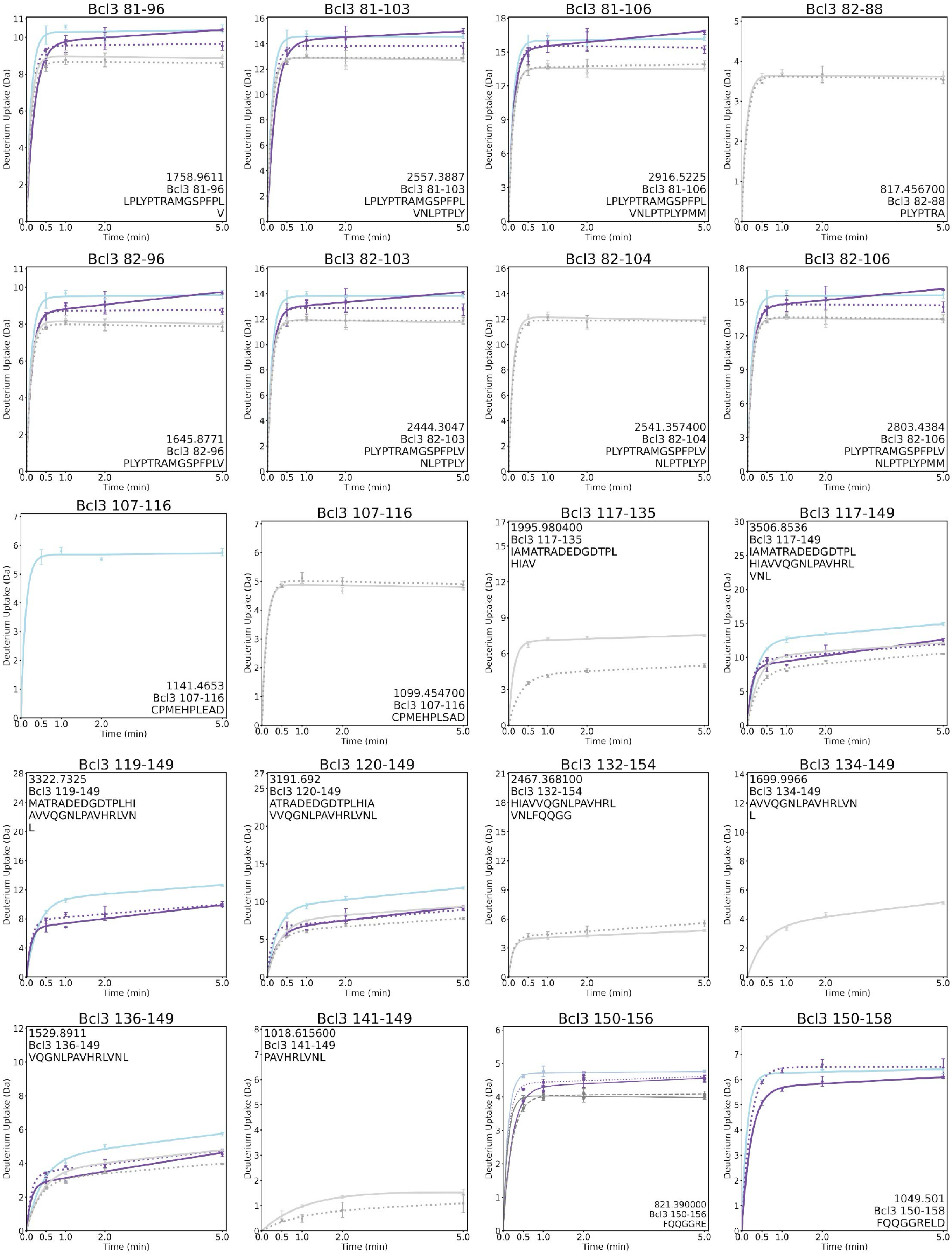
Bcl3 81-96
1758.9611
Bcl3 81-96
LPLYPTRAMGSPFPL
V
Bcl3 81-103
2557.3887
Bcl3 81-103
LPLYPTRAMGSPFPL
VNLPTPLY
Bcl3 81-106
2916.5225
Bcl3 81-106
LPLYPTRAMGSPFPL
VNLPTPLYPMM
Bcl3 82-88
817.456700
Bcl3 82-88
PLYPTRA
Bcl3 82-96
1645.8771
Bcl3 82-96
PLYPTRAMGSPFPLV
Bcl3 82-103
2444.3047
Bcl3 82-103
PLYPTRAMGSPFPLV
NLPTPLY
Bcl3 82-104
2541.357400
Bcl3 82-104
PLYPTRAMGSPFPLV
NLPTPLYP
Bcl3 82-106
2803.4384
Bcl3 82-106
PLYPTRAMGSPFPLV
NLPTPLYPMM
Bcl3 107-116
1141.4653
Bcl3 107-116
CPMEHPLEAD
Bcl3 107-116
1099.454700
Bcl3 107-116
CPMEHPLSAD
Bcl3 117-135
1995.980400
Bcl3 117-135
IAMATRADEDGDTPL
HIAV
Bcl3 117-149
3506.8536
Bcl3 117-149
IAMATRADEDGDTPL
HIAVVQGNLPAVHRL
VNL
Bcl3 119-149
3322.7325
Bcl3 119-149
MATRADEDGDTPLHI
AVVQGNLPAVHRLVN
L
Bcl3 120-149
3191.692
Bcl3 120-149
ATRADEDGDTPLHIA
VVQGNLPAVHRLVNL
Bcl3 132-154
2467.368100
Bcl3 132-154
HIAVVQGNLPAVHRL
VNLFQQGG
Bcl3 134-149
1699.9966
Bcl3 134-149
AVVQGNLPAVHRLVN
L
Bcl3 136-149
1529.8911
Bcl3 136-149
VQGNLPAVHRLVNL
Bcl3 141-149
1018.615600
Bcl3 141-149
PAVHRLVNL
Bcl3 150-156
821.390000
Bcl3 150-156
FQQGGRE
Bcl3 150-158
1049.501
Bcl3 150-158
FQQGGRELD
Deuterium Uptake (Da)
Time (min)

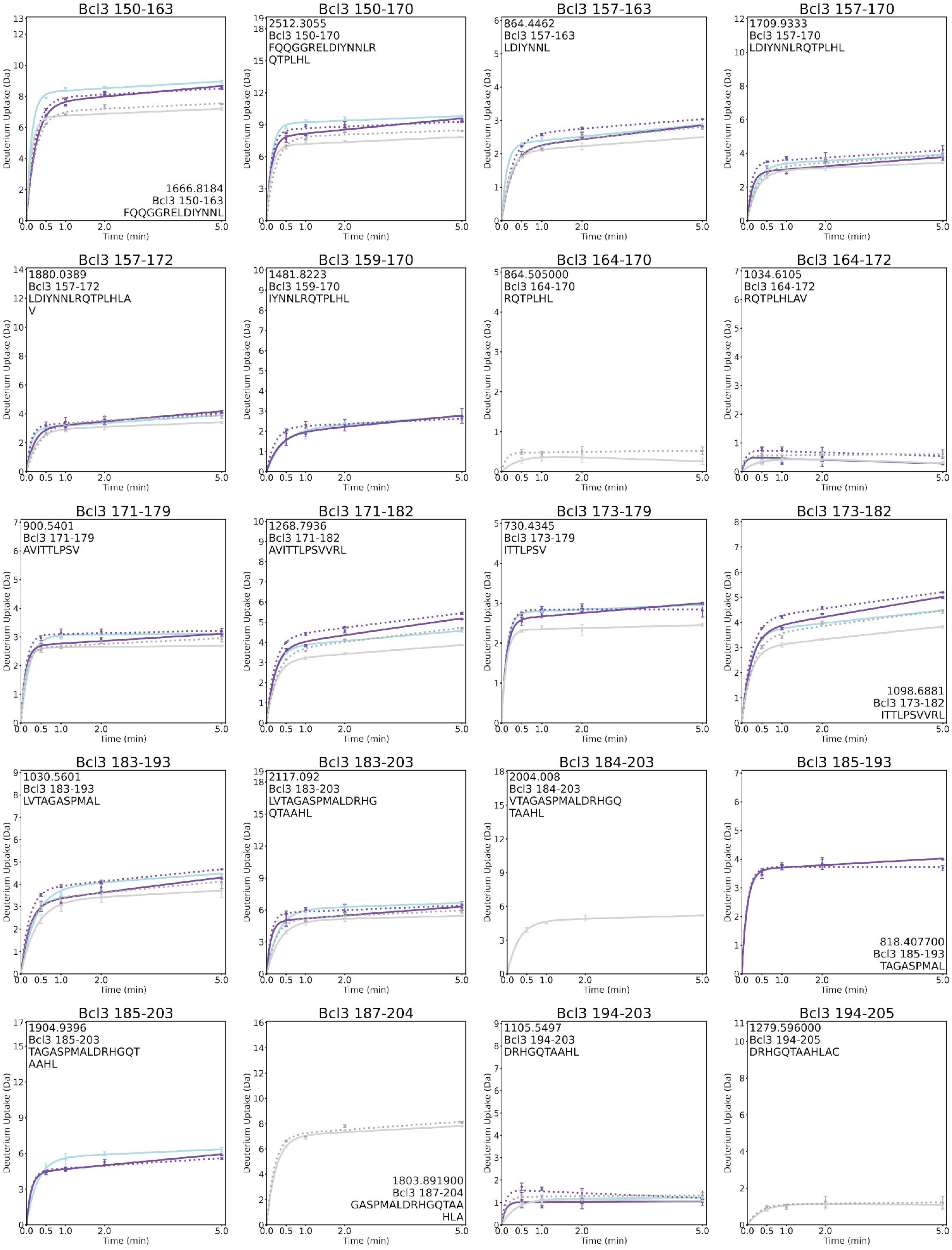
Bcl3 150-163
1666.8184
Bcl3 150-163
FQQGGRELDIYNNL
Bcl3 150-170
2512.3055
Bcl3 150-170
FQQGGRELDIYNNLR
QTPLHL
Bcl3 157-163
864.4462
Bcl3 157-163
LDIYNNL
Bcl3 157-170
1709.9333
Bcl3 157-170
LDIYNNLRQTPLHL
Bcl3 157-172
1880.0389
Bcl3 157-172
LDIYNNLRQTPLHLA
V
Bcl3 159-170
1481.8223
Bcl3 159-170
IYNNLRQTPLHL
Bcl3 164-170
864.505000
Bcl3 164-170
RQTPLHL
Bcl3 164-172
1034.6105
Bcl3 164-172
RQTPLHLAV
Bcl3 171-179
900.5401
Bcl3 171-179
AVITTLPSV
Bcl3 171-182
1268.7936
Bcl3 171-182
AVITTLPSVVRL
Bcl3 173-179
730.4345
Bcl3 173-179
ITTLPSV
Bcl3 173-182
1098.6881
Bcl3 173-182
ITTLPSVVRL
Bcl3 183-193
1030.5601
Bcl3 183-193
LVTAGASPMAL
Bcl3 183-203
2117.092
Bcl3 183-203
LVTAGASPMALDRHG
QTAAHL
Bcl3 184-203
2004.008
Bcl3 184-203
VTAGASPMALDRHGQ
TAAHL
Bcl3 185-193
818.407700
Bcl3 185-193
TAGASPMAL
Bcl3 185-203
1904.9396
Bcl3 185-203
TAGASPMALDRHGQT
AAHL
Bcl3 187-204
1803.891900
Bcl3 187-204
GASPMALDRHGQTAA
HLA
Bcl3 194-203
1105.5497
Bcl3 194-203
DRHGQTAAHL
Bcl3 194-205
1279.596000
Bcl3 194-205
DRHGQTAAHLAC
Deuterium Uptake (Da)
Time (min)

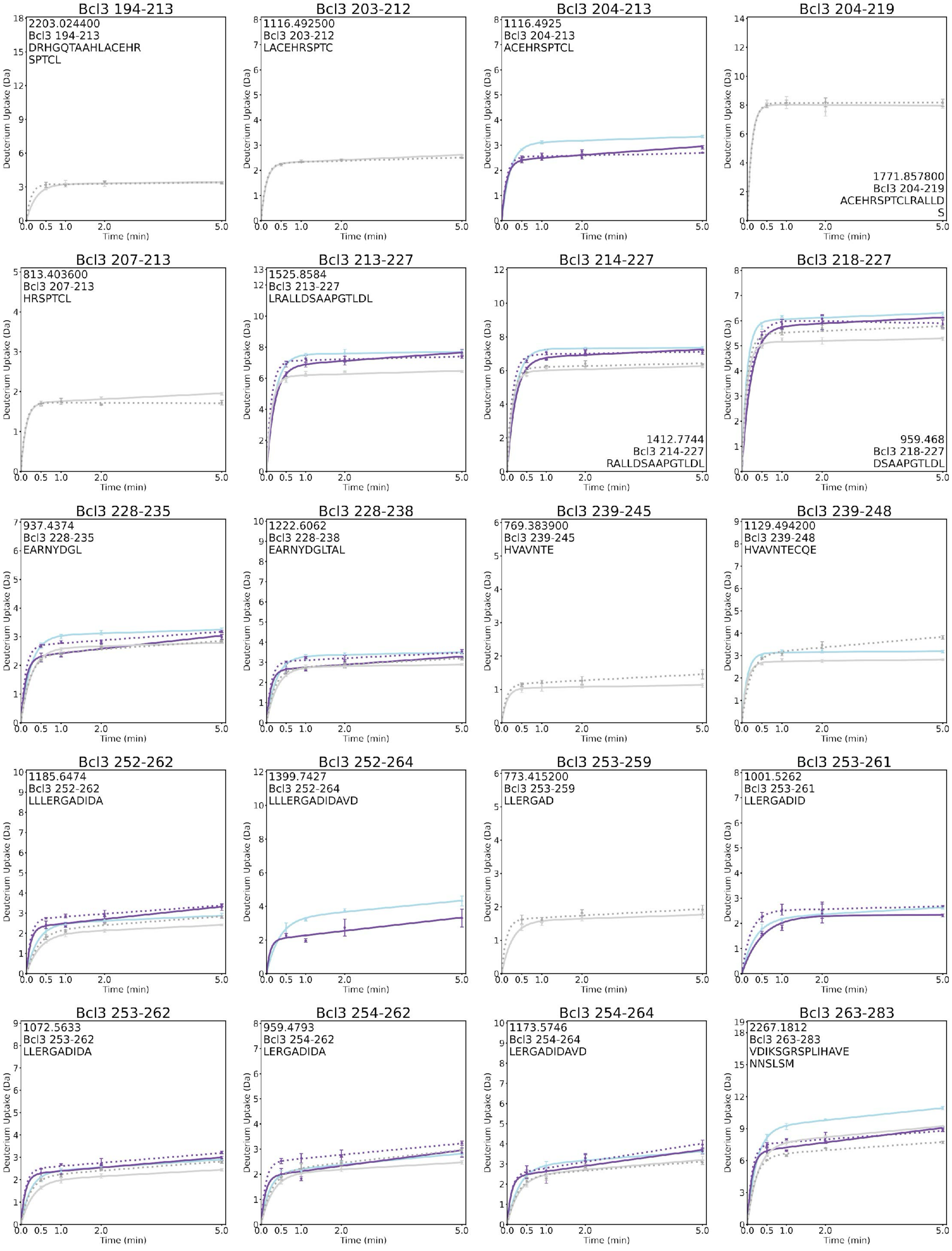
Bcl3 194-213
2203.024400
Bcl3 194-213
DRHGQTAAHLACEHR
SPTCL
Bcl3 203-212
1116.492500
Bcl3 203-212
LACEHRSPTC
Bcl3 204-213
1116.4925
Bcl3 204-213
ACEHRSPTCL
Bcl3 204-219
1771.857800
Bcl3 204-219
ACEHRSPTCLRALLD
S
Bcl3 207-213
813.403600
Bcl3 207-213
HRSPTCL
Bcl3 213-227
1525.8584
Bcl3 213-227
LRALLDSAAPGTLDL
Bcl3 214-227
1412.7744
Bcl3 214-227
RALLDSAAPGTLDL
Bcl3 218-227
959.468
Bcl3 218-227
DSAAPGTLDL
Bcl3 228-235
937.4374
Bcl3 228-235
EARNYDGL
Bcl3 228-238
1222.6062
Bcl3 228-238
EARNYDGLTAL
Bcl3 239-245
769.383900
Bcl3 239-245
HVAVNTE
Bcl3 239-248
1129.494200
Bcl3 239-248
HVAVNTECQE
Bcl3 252-262
1185.6474
Bcl3 252-262
LLLERGADIDA
Bcl3 252-264
1399.7427
Bcl3 252-264
LLLERGADIDAVD
Bcl3 253-259
773.415200
Bcl3 253-259
LLERGAD
Bcl3 253-261
1001.5262
Bcl3 253-261
LLERGADID
Bcl3 253-262
1072.5633
Bcl3 253-262
LLERGADIDA
Bcl3 254-262
959.4793
Bcl3 254-262
LERGADIDA
Bcl3 254-264
1173.5746
Bcl3 254-264
LERGADIDAVD
Bcl3 263-283
2267.1812
Bcl3 263-283
VDIKSGRSPLIHAVE
NNSLSM
Deuterium Uptake (Da)
Time (min)

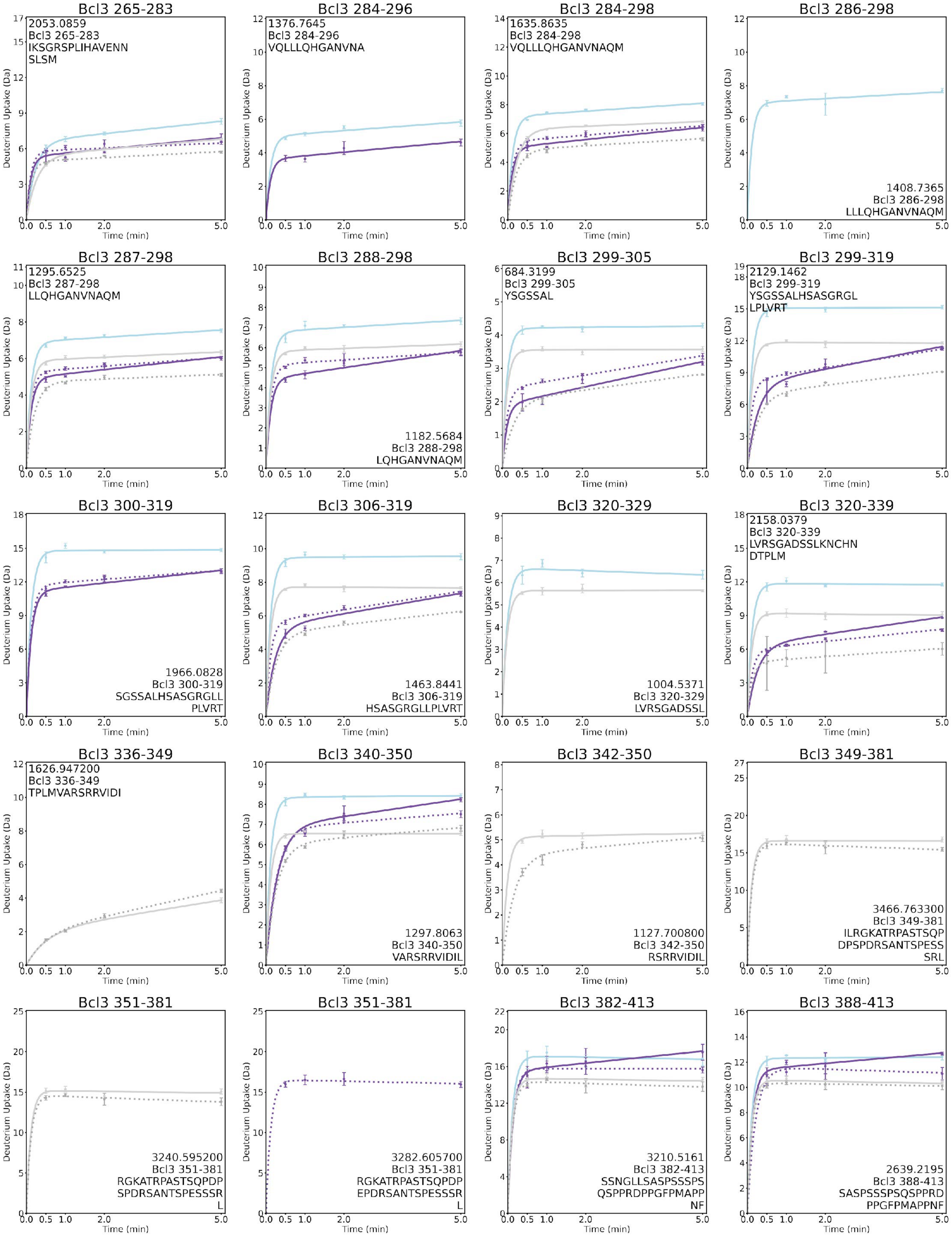
Bcl3 265-283
2053.0859
Bcl3 265-283
IKSGRSPLIHAVENN
SLSM
Bcl3 284-296
1376.7645
Bcl3 284-296
VQLLLQHGANVNA
Bcl3 284-298
1635.8635
Bcl3 284-298
VQLLLQHGANVNAQM
Bcl3 286-298
1408.7365
Bcl3 286-298
LLLQHGANVNAQM
Bcl3 287-298
1295.6525
Bcl3 287-298
LLQHGANVNAQM
Bcl3 288-298
1182.5684
Bcl3 288-298
LQHGANVNAQM
Bcl3 299-305
684.3199
Bcl3 299-305
YSGSSAL
Bcl3 299-319
2129.1462
Bcl3 299-319
YSGSSALHSASGRGL
LPLVRT
Bcl3 300-319
1966.0828
Bcl3 300-319
SGSSALHSASGRGLL
PLVRT
Bcl3 306-319
1463.8441
Bcl3 306-319
HSASGRGLLPLVRT
Bcl3 320-329
1004.5371
Bcl3 320-329
LVRSGADSSL
Bcl3 320-339
2158.0379
Bcl3 320-339
LVRSGADSSLKNCHN
DTPLM
Bcl3 336-349
1626.947200
Bcl3 336-349
TPLMVARSRRVIDI
Bcl3 340-350
1297.8063
Bcl3 340-350
VARSRRVIDIL
Bcl3 342-350
1127.700800
Bcl3 342-350
RSRRVIDIL
Bcl3 349-381
3466.763300
Bcl3 349-381
ILRGKATRPASTSQP
DPSPDRSANTSPESS
SRL
Bcl3 351-381
3240.595200
Bcl3 351-381
RGKATRPASTSQPDP
SPDRSANTSPESSSR
L
Bcl3 351-381
3282.605700
Bcl3 351-381
RGKATRPASTSQPDP
EPDRSANTSPESSSR
L
Bcl3 382-413
3210.5161
Bcl3 382-413
SSNGLLSASPSSSPS
QSPPRDPPGFPMAPP
NF
Bcl3 388-413
2639.2195
Bcl3 388-413
SASPSSSPSQSPPRD
PPGFPMAPPNF
Deuterium Uptake (Da)
Time (min)

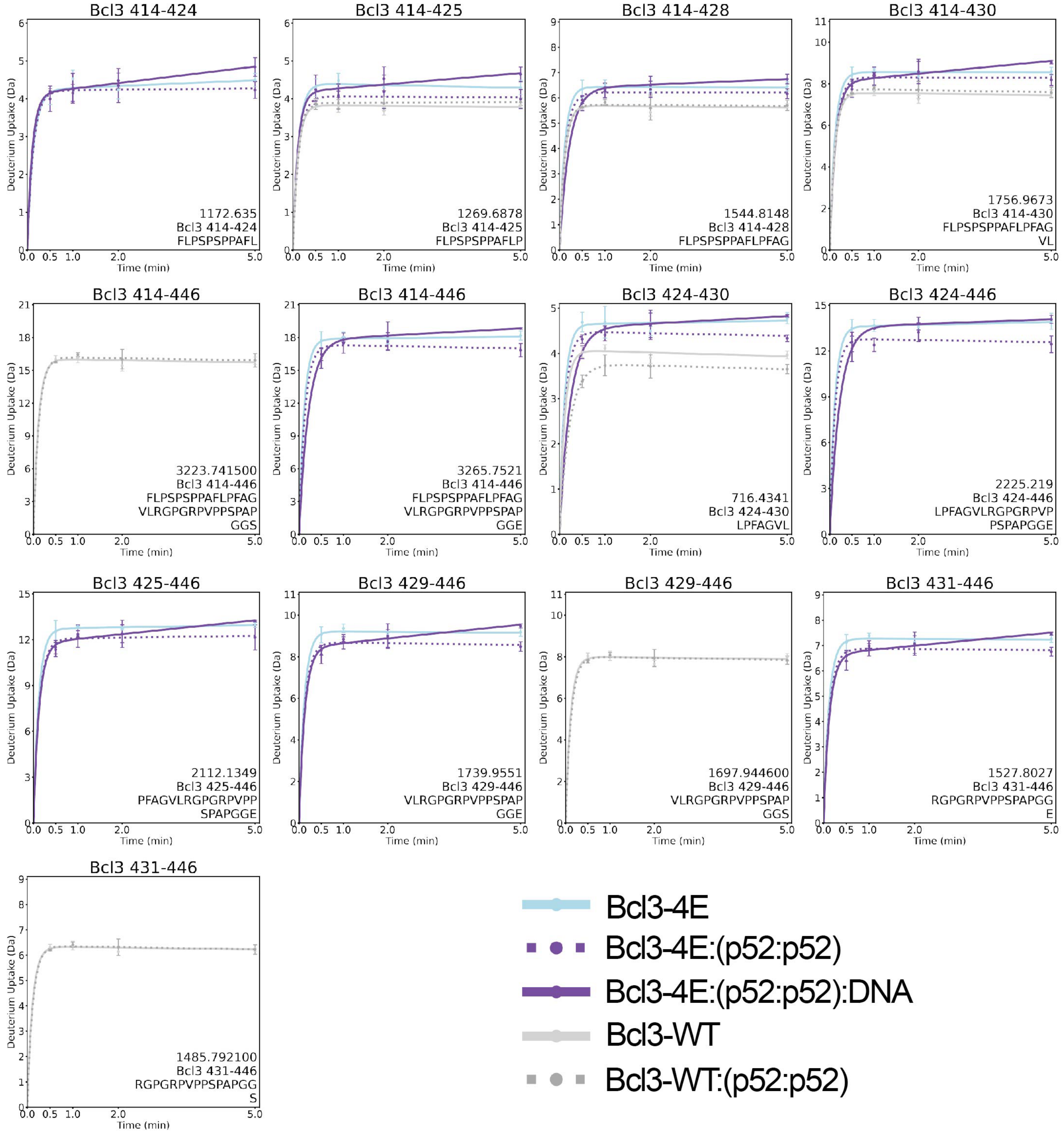


**Figure S2. Deuterium uptake plots of Bcl3 peptides.** HDX-MS uptake plots of Bcl3-WT and Bcl3-4E peptides showing deuterium uptakes at 0.5, 1, 2, and 5 minute time-points within Bcl3-WT (grey), Bcl3-WT:(p52:p52) (grey-dot), Bcl3-4E (blue), Bcl3-4E:(p52:p52) (violet-dot) and Bcl3-4E:(p52:p52):DNA (violet) complexes. All analyses were performed in experimental triplicate and error bars are shown.

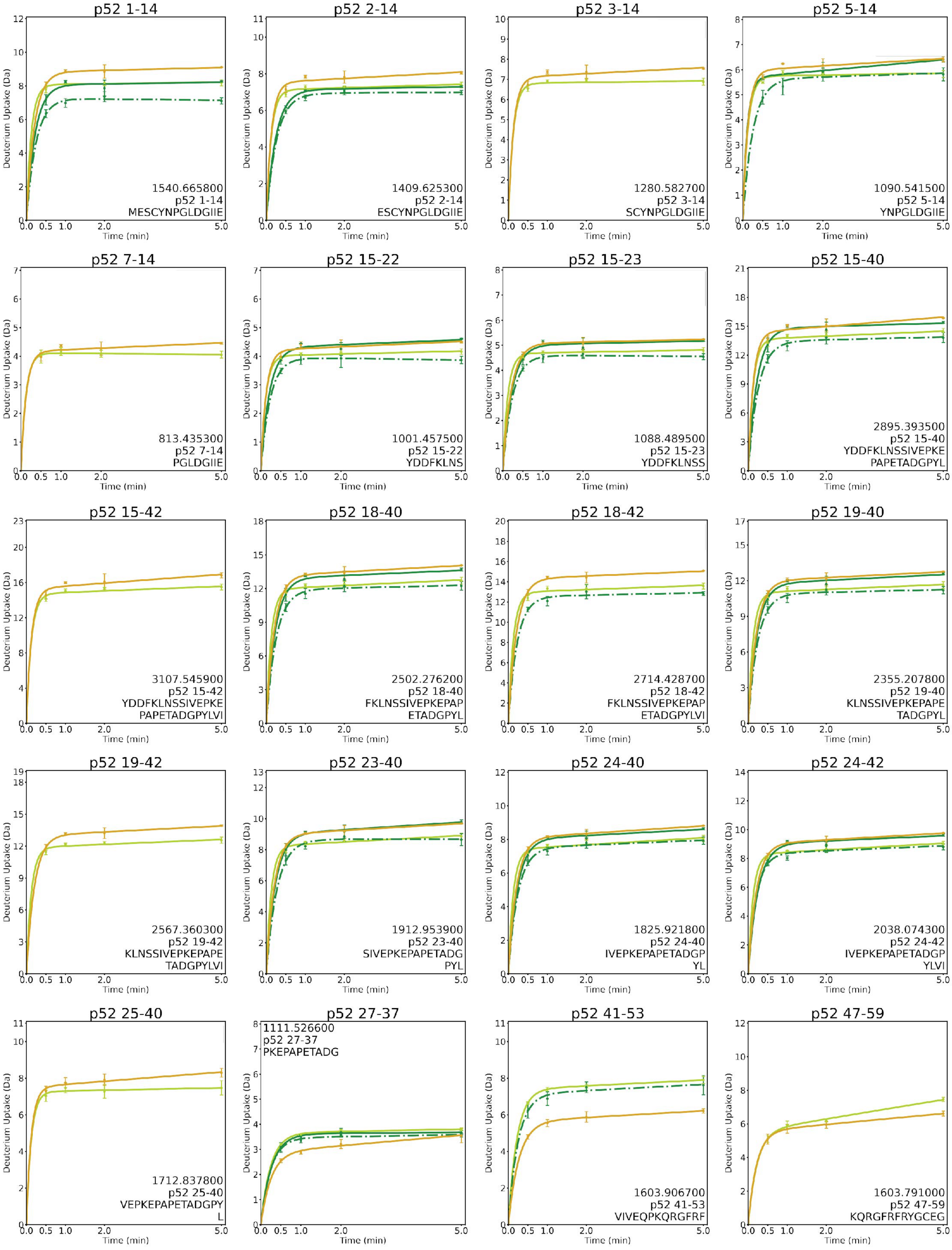
Deuterium Uptake (Da)
Time (min)
p52 1-14
1540.665800
p52 1-14
MESCYNPGLDGIIE
p52 2-14
1409.625300
p52 2-14
ESCYNPGLDGIIE
p52 3-14
1280.582700
p52 3-14
SCYNPGLDGIIE
p52 5-14
1090.541500
p52 5-14
YNPGLDGIIE
p52 7-14
813.435300
p52 7-14
PGLDGIIE
p52 15-22
1001.457500
p52 15-22
YDDFKLNS
p52 15-23
1088.489500
p52 15-23
YDDFKLNSS
p52 15-40
2895.393500
p52 15-40
YDDFKLNSSIVEPKE
PAPETADGPYL
p52 15-42
3107.545900
p52 15-42
YDDFKLNSSIVEPKE
PAPETADGPYLVI
p52 18-40
2502.276200
p52 18-40
FKLNSSIVEPKEPAP
ETADGPYL
p52 18-42
2714.428700
p52 18-42
FKLNSSIVEPKEPAP
ETADGPYLVI
p52 19-40
2355.207800
p52 19-40
KLNSSIVEPKEPAPE
TADGPYL
p52 19-42
2567.360300
p52 19-42
KLNSSIVEPKEPAPE
TADGPYLVI
p52 23-40
1912.953900
p52 23-40
SIVEPKEPAPETADG
PYL
p52 24-40
1825.921800
p52 24-40
IVEPKEPAPETADGP
YL
p52 24-42
2038.074300
p52 24-42
IVEPKEPAPETADGP
YLVI
p52 25-40
1712.837800
p52 25-40
VEPKEPAPETADGPY
L
p52 27-37
1111.526600
p52 27-37
PKEPAPETADG
p52 41-53
1603.906700
p52 41-53
VIVEQPKQRGFRF
p52 47-59
1603.791000
p52 47-59
KQRGFRFRYGCEG

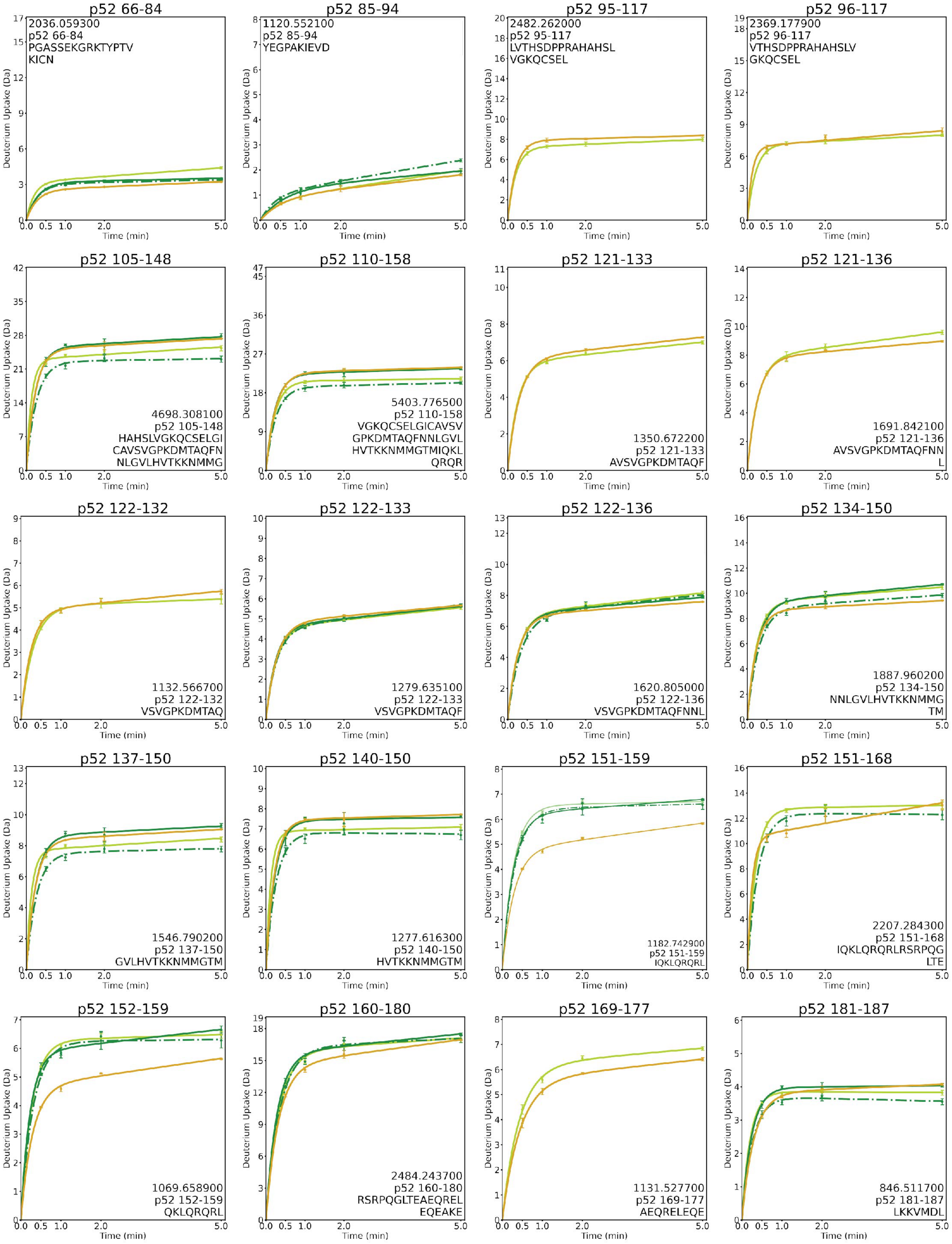
p52 66-84
2036.059300
p52 66-84
PGASSEKGRKTYPTV
KICN
p52 85-94
1120.552100
p52 85-94
YEGPAKIEVD
p52 95-117
2482.262000
p52 95-117
LVTHSDPPRAHAHSL
VGKQCSEL
p52 96-117
2369.177900
p52 96-117
VTHSDPPRAHAHSLV
GKQCSEL
p52 105-148
4698.308100
p52 105-148
HAHSLVGKQCSELGI
CAVSVGPKDMTAQFN
NLGVLHVTKKNMMG
p52 110-158
5403.776500
p52 110-158
VGKQCSELGICAVSV
GPKDMTAQFNNLGVL
HVTKKNMMGTMIQKL
QRQR
p52 121-133
1350.672200
p52 121-133
AVSVGPKDMTAQF
p52 121-136
1691.842100
p52 121-136
AVSVGPKDMTAQFNN
L
p52 122-132
1132.566700
p52 122-132
VSVGPKDMTAQ
p52 122-133
1279.635100
p52 122-133
VSVGPKDMTAQF
p52 122-136
1620.805000
p52 122-136
VSVGPKDMTAQFNNL
p52 134-150
1887.960200
p52 134-150
NNLGVLHVTKKNMMG
TM
p52 137-150
1546.790200
p52 137-150
GVLHVTKKNMMGTM
p52 140-150
1277.616300
p52 140-150
HVTKKNMMGTM
p52 151-159
1182.742900
p52 151-159
IQKLQRQRL
p52 151-168
2207.284300
p52 151-168
IQKLQRQRLRSRPQG
LTE
p52 152-159
1069.658900
p52 152-159
QKLQRQRL
p52 160-180
2484.243700
p52 160-180
RSRPQGLTEAEQREL
EQEAKE
p52 169-177
1131.527700
p52 169-177
AEQRELEQE
p52 181-187
846.511700
p52 181-187
LKKVMDL
Deuterium Uptake (Da)
Time (min)

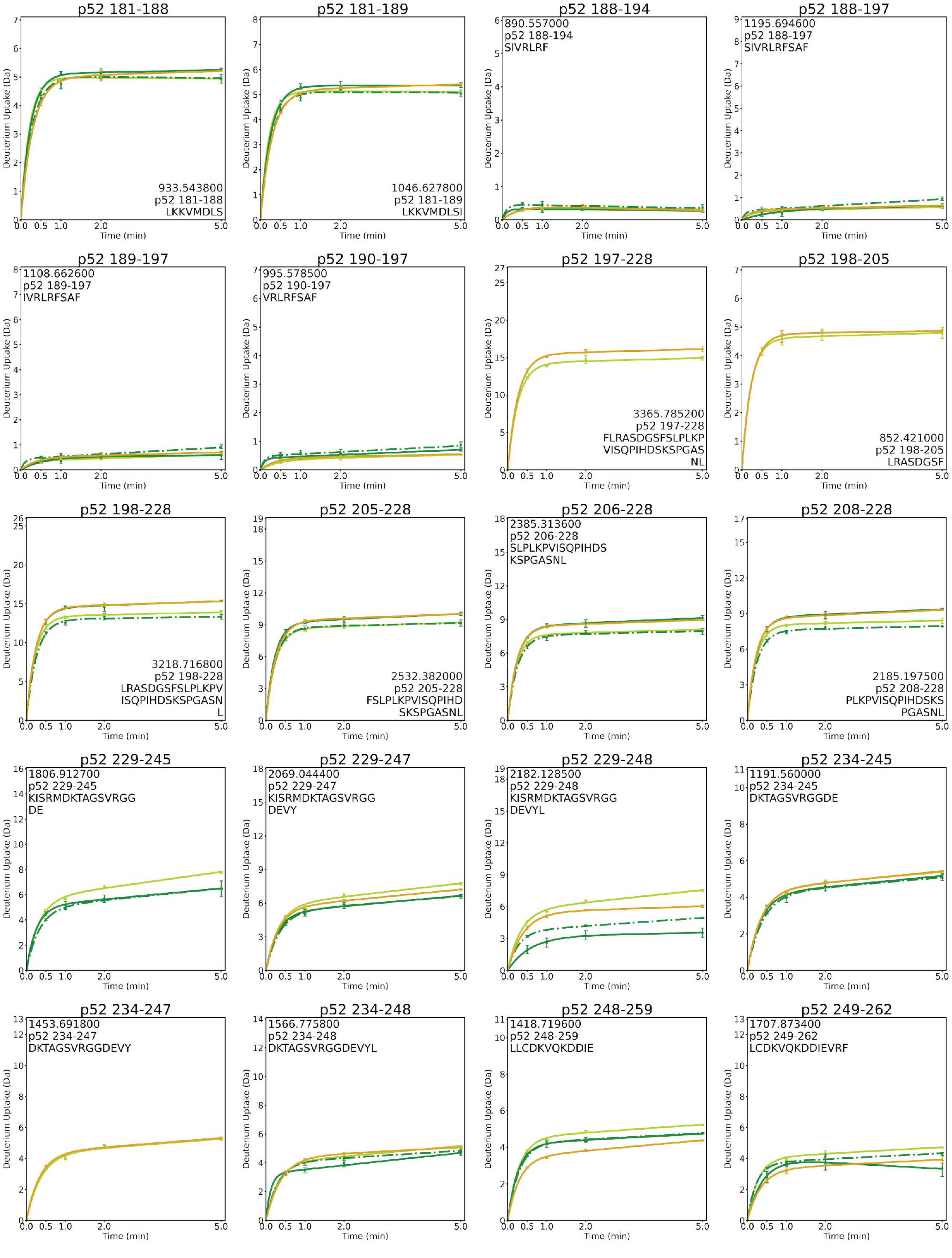
p52 181-188
933.543800
p52 181-188
LKKVMDLS
p52 181-189
1046.627800
p52 181-189
LKKVMDLSI
p52 188-194
890.557000
p52 188-194
SIVRLRF
p52 188-197
1195.694600
p52 188-197
SIVRLRFSAF
p52 189-197
1108.662600
p52 189-197
IVRLRFSAF
p52 190-197
995.578500
p52 190-197
VRLRFSAF
p52 197-228
3365.785200
p52 197-228
FLRASDGSFSLPLKP
VISQPIHDSKSPGAS
NL
p52 198-205
852.421000
p52 198-205
LRASDGSF
p52 198-228
3218.716800
p52 198-228
LRASDGSFSLPLKPV
ISQPIHDSKSPGASN
L
p52 205-228
2532.382000
p52 205-228
FSLPLKPVISQPIHD
SKSPGASNL
p52 206-228
2385.313600
p52 206-228
SLPLKPVISQPIHDS
KSPGASNL
p52 208-228
2185.197500
p52 208-228
PLKPVISQPIHDSKS
PGASNL
p52 229-245
1806.912700
p52 229-245
KISRMDKTAGSVRGG
DE
p52 229-247
2069.044400
p52 229-247
KISRMDKTAGSVRGG
DEVY
p52 229-248
2182.128500
p52 229-248
KISRMDKTAGSVRGG
DEVYL
p52 234-245
1191.560000
p52 234-245
DKTAGSVRGGDE
p52 234-247
1453.691800
p52 234-247
DKTAGSVRGGDEVY
p52 234-248
1566.775800
p52 234-248
DKTAGSVRGGDEVYL
p52 248-259
1418.719600
p52 248-259
LLCDKVQKDDIE
p52 249-262
1707.873400
p52 249-262
LCDKVQKDDIEVRF
Deuterium Uptake (Da)
Time (min)

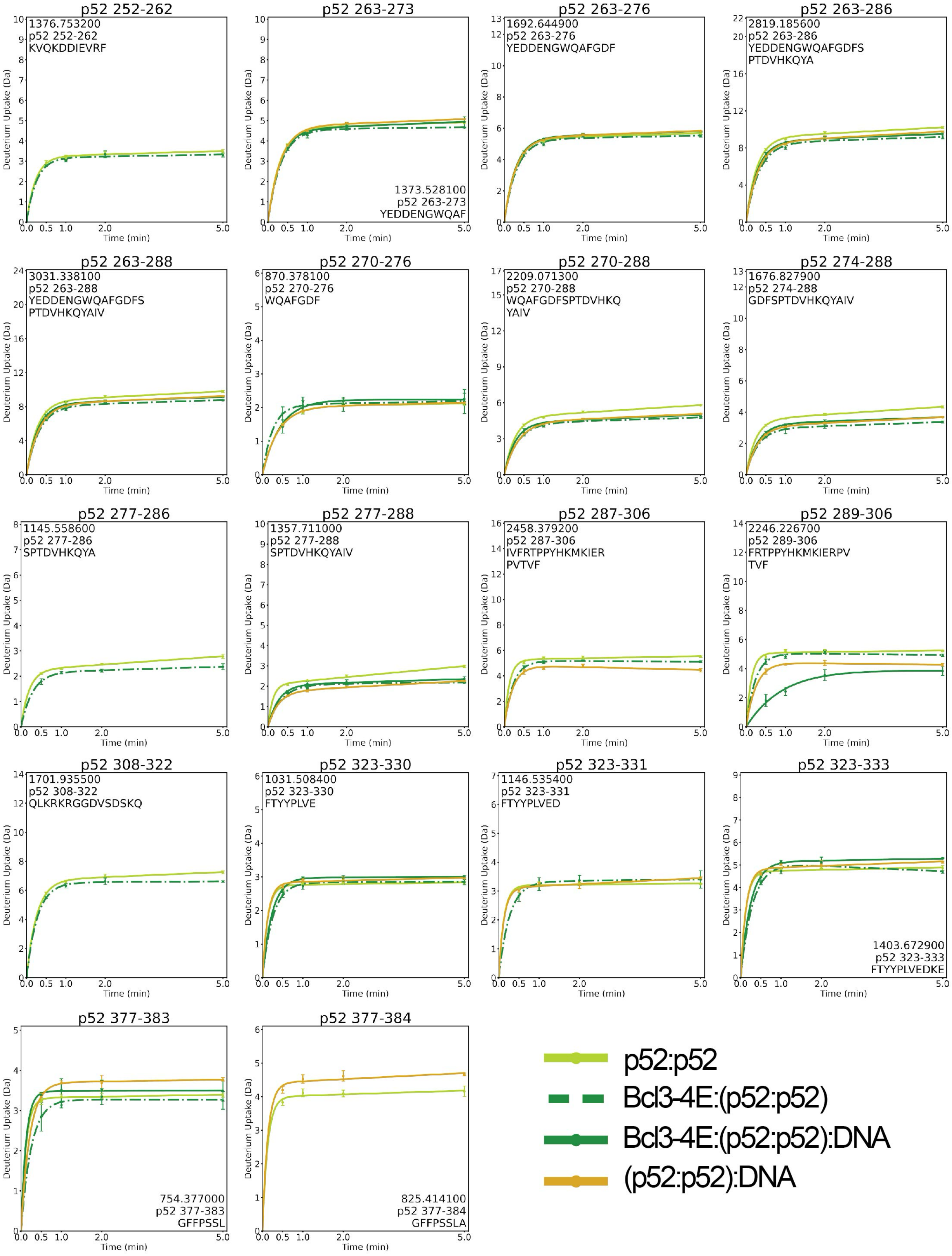
p52 252-262
1376.753200
p52 252-262
KVQKDDIEVRF
p52 263-273
1373.528100
p52 263-273
YEDDENGWQAF
p52 263-276
1692.644900
p52 263-276
YEDDENGWQAFGDF
p52 263-286
2819.185600
p52 263-286
YEDDENGWQAFGDFS
PTDVHKQYA
p52 263-288
3031.338100
p52 263-288
YEDDENGWQAFGDFS
PTDVHKQYAIV
p52 270-276
870.378100
p52 270-276
WQAFGDF
p52 270-288
2209.071300
p52 270-288
WQAFGDFSPTDVHKQ
YAIV
p52 274-288
1676.827900
p52 274-288
GDFSPTDVHKQYAIV
p52 277-286
1145.558600
p52 277-286
SPTDVHKQYA
p52 277-288
1357.711000
p52 277-288
SPTDVHKQYAIV
p52 287-306
2458.379200
p52 287-306
IVFRTPPYHKMKIER
PVTVF
p52 289-306
2246.226700
p52 289-306
FRTPPYHKMKIERPV
TVF
p52 308-322
1701.935500
p52 308-322
QLKRKRGGDVSDSKQ
p52 323-330
1031.508400
p52 323-330
FTYYPLVE
p52 323-331
1146.535400
p52 323-331
FTYYPLVED
p52 323-333
1403.672900
p52 323-333
FTYYPLVEDKE
p52 377-383
754.377000
p52 377-383
GFFPSSL
p52 377-384
825.414100
p52 377-384
GFFPSSLA
Deuterium Uptake (Da)
Time (min)
p52:p52
Bcl3-4E:(p52:p52)
Bcl3-4E:(p52:p52):DNA
(p52:p52):DNA

**Figure S3. Deuterium uptake plots of p52 peptides.** HDX-MS uptake plots of p52 peptides showing deuterium uptakes at 0.5, 1, 2, and 5 minute time-points within p52:p52 homodimer (light green), Bcl3-4E:(p52:p52) (dark green dash-dot), Bcl3-4E:(p52:p52):DNA (dark green) and (p52:p52):DNA (golden) complexes. All analyses were performed in experimental triplicate and error bars are shown.

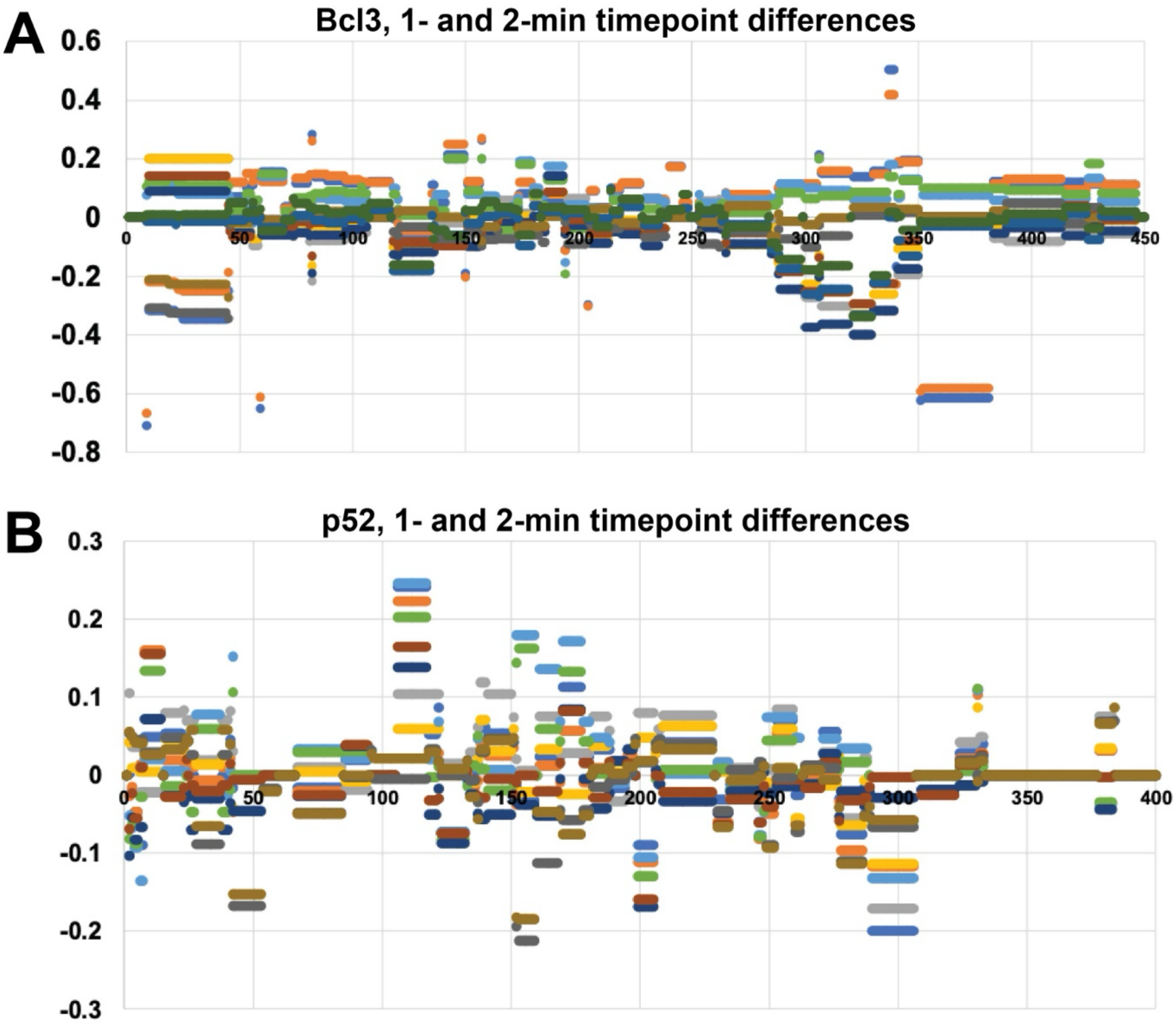


**Figure S4. Difference in fractional uptake.** Plots displaying spread of 'difference in fractional uptake' at 1- and 2-minute timepoint of (A) Bcl3 and (B) p52 peptides within different complexes.

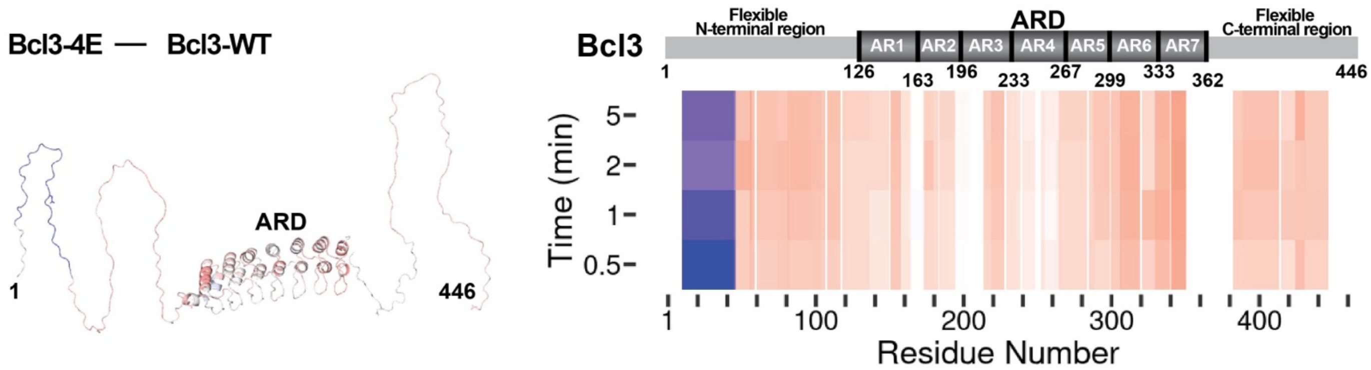


**Figure S5. Heatmaps showing the difference in fractional uptake of deuterium in Bcl3-4E vs. Bcl3-WT in as free proteins.**

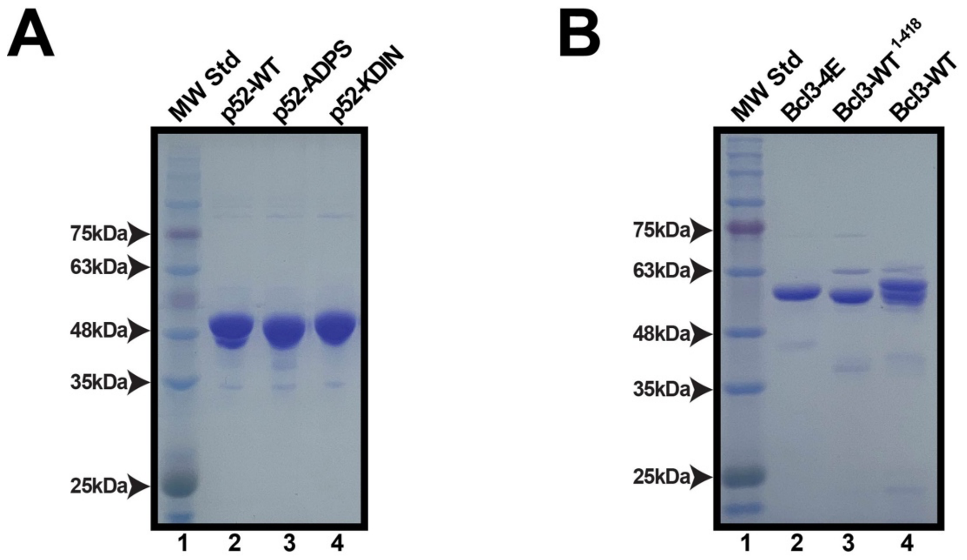


**Figure S6. Purity of recombinant proteins.** (A) SDS-PAGE analysis showing the p52-WT, p52-ADPS and -KDIN mutants in complex with phospho-mimetic Bcl3-4E with similar purity. (B) SDS-PAGE analysis showing the His-Bcl3-4E, His-Bcl3-WT$^{1\text{-}418}$, and His-Bcl3-WT with similar purity.
